# Exploiting the Achilles’ Heel of Viral RNA Processing to Develop Novel Antivirals

**DOI:** 10.1101/2024.09.11.612277

**Authors:** Ali Zahedi Amiri, Choudhary Ahmed, Subha Dahal, Filomena Grosso, Haomin Leng, Peter Stoilov, Maria Mangos, Johanne Toutant, Lulzim Shkreta, Liliana Attisano, Benoit Chabot, Martha Brown, Mario Huesca, Alan Cochrane

## Abstract

Viruses continue to pose a significant health burden to the human population, and recent history has shown a concerning surge in viral threats. Treatment options for viral infections are limited, and viruses have proven adept at evolving resistance to many existing therapies, highlighting a significant vulnerability in our defenses. In response to this challenge, we explored the modulation of cellular RNA metabolic processes as an alternative paradigm to antiviral development. Many viruses depend on the host cell’s RNA splicing machinery, and small alterations to this host process results in catastrophic changes in viral protein production, ultimately inhibiting virus replication. Previously, the small molecule 5342191 was identified as a potent inhibitor of HIV-1 replication by altering viral RNA accumulation at doses that minimally affect host gene expression. In this report, we document 5342191 as a potent inhibitor of adenovirus, coronavirus, and influenza replication. In each case, 5342191-mediated reduction in virus replication was associated with altered viral RNA accumulation and loss of viral structural protein expression. Interestingly, while resistant viruses were rapidly isolated for compounds targeting either virus-encoded proteases or polymerases, we have not yet isolated 534219-resistant variants of coronavirus or influenza. As with HIV-1, 5342191’s inhibition of coronaviruses and influenza is mediated through the activation of specific cell signaling networks, including GPCR and/or MAPK signaling pathways that ultimately affect SR kinase expression. Together, these studies highlight the therapeutic potential of compounds that target cellular processes essential for the replication of multiple viruses. Not only do these compounds hold promise as broad-spectrum antivirals, but they also offer the potential of greater resilience in combating viral infections.

## Introduction

Over the past several decades, humans have experienced multiple viral epidemics and pandemics, including HIV-1, coronaviruses (SARS, MERS, SARS-CoV-2), influenza (H1N1), Zika, and Ebola, each causing significant morbidity and mortality^1^. To avoid the social and fiscal upheaval experienced during the most recent COVID-19 pandemic, efforts must be directed at preparing for multiple threats simultaneously. Although vaccines can be effective at reducing the impact of an infection, their high specificity and the need for administration several weeks to months prior to exposure limit their utility for those already infected. In some instances, vaccines can even enhance pathology, as seen with Dengue and Zika^2,3^. Furthermore, in several instances, despite significant effort, development of effective vaccines has taken decades (e.g. respiratory syncytial virus (RSV), Dengue) or has not occurred at all (e.g., HIV-1, Hepatitis C virus (HCV)).

Small molecule inhibitors of virus replication are a key, complementary approach to vaccines in mitigating the immediate and long-term impact of a new pandemic. These antiviral compounds reduce disease symptoms in infected individuals or can be used to prevent infection. In contrast to vaccines, antivirals are active within hours to days of administration, facilitating a more rapid response to an emerging pandemic or epidemic. Hence, the availability of compounds with proven activity against multiple virus families and their prequalification for use in humans is essential for pandemic preparedness against existing or emerging viral threats. However, of the 220 known human viruses, only 10 have small molecule inhibitors^1^.

Development of antivirals has typically focused on inhibitors of viral-encoded proteins such as the viral polymerase or protease^1,4^. While successful, the utility of these drugs, termed direct-acting antivirals (DAAs), against unrelated viruses is highly variable, and rapid virus evolution can quickly diminish drug efficacy due to the emergence of resistant strains^1^. The modulation of cellular processes essential for virus replication (host-directed therapies, HDT) is an alternative treatment modality^5–8^. As obligate pathogens, viruses depend upon multiple host cellular processes for genome replication/expression and new virion assembly/release. Since multiple viruses can depend on the same cellular processes, altering the accessibility or activity of these cellular processes can impact the replication of multiple unrelated viruses. Even small alterations in key host cellular processes can suppress viral reproduction to prevent pathology, giving the immune system time to control the virus. This allows effective dosages of HDT antivirals to be used, even if they have known toxicities at higher concentrations. Furthermore, in several examples of HDT antivirals, the emergence of resistant virus strains has been slow or nonexistent due to the inability of the virus to compensate for the lost cellular function^9^. Target processes for such HDTs include lipid metabolism, cell signaling, membrane trafficking, protein quality control, and RNA processing ^8,10^.

Previous studies by our group have focused on the modulation of HIV-1 RNA processing by small molecules to inhibit its replication. Recently, we demonstrated that three HIV-1 inhibitors GPS491, harmine, and cardiotonic steroids-also have the capacity to inhibit the replication of several unrelated viruses (including adenovirus and coronaviruses like SARS-CoV-2) by affecting the production and processing of their respective RNAs^11–13^. To expand the inventory of HDTs, we examined the capacity of 5342191 (N-[4-chloro-3-(trifluoromethyl)phenyl]-7-nitro-2,1,3-benzoxadiazol-4-amine), another inhibitor of HIV-1 identified by our group^14^, to inhibit the replication of adenovirus, coronaviruses, and influenza. While the dose of 5342191 required to fully suppress virus replication differed among the viruses tested, the compound inhibited replication of these viruses at doses well below those affecting cell viability. For each virus, inhibition was associated with changes in viral RNA abundance and/or processing at concentrations at or below those previously shown to have minimal effects on host gene expression or RNA splicing ^14^.

Similar to the response seen for HIV-1, the antiviral effect of 5342191 against both HCoV-229E and influenza PR8, was reversed upon treatment of cells with inhibitors of the MAPK or GPCR-dependent signaling pathways. Furthermore, attempts to generate 5342191-resistant variants of either HCoV-229E and influenza PR8 met with limited success after 10 or 4 rounds of selection, respectively. In contrast, resistant forms of both HCoV-229E and influenza PR8 to two DAAs were isolated after only 5 rounds of selection. Together, these findings establish 5342191 as a broad spectrum antiviral capable of targeting several viruses of concern by affecting the accumulation and/or processing of viral RNAs, adding to the inventory of compounds that could be developed to respond to future virus outbreaks.

## 2 , Materials & Methods

### 2.1 Cell lines and virus strains

A549 cells (human lung carcinoma) were obtained from the American Type Culture Collection (ATCC) at passage level 76 and used between passages 89 and 110. HEK 293 (human embryonic kidney) cells were obtained from F. Graham, McMaster University, Hamilton, Ontario, Canada, at passage 24 and were used between passages 58 and 90. A549 and HEK 293 cells were cultured in Eagle’s minimum essential medium (MEM, Gibco) supplemented with 10% fetal bovine serum (FBS, Sigma-Aldrich) plus penicillin (100 U/mL) and streptomycin (100 μg/ml). Huh7 cells for infection with coronaviruses were maintained in Dulbecco’s Modified Eagle Medium (DMEM) with high glucose (HyClone) supplemented with 10% heat-inactivated FBS and 1% penicillin/streptomycin. All the cell lines were maintained at 37°C in a humidified incubator with 5% CO_2_. Adenovirus used in the studies was HAdV-C5. Coronaviruses 229E, OC43, and SB2, Delta, and Omicron BA.1 variants of SARS-CoV2 were obtained from Dr. S. Mubareka (University of Toronto, Toronto, ON, Canada) and the combined Containment Level 3 unit at the University of Toronto, Toronto, ON, Canada. Influenza PR8 (A/PR/8/34 (H1N1; PR8) and MDCK cells (Madin-Darby Canine Kidney cells) were obtained from Dr. Tania Watts, Dept. of Immunology, University of Toronto.

### 2.2 Generation of human lung organoids (hLOs)

Human lung organoids were generated in the Applied Organoid Core (ApOC) Facility (Donnelly Centre, University of Toronto) with a modified protocol using H9 human embryonic stem cells (obtained from the WiCell Research Institute) and 30 day old Air-liquid Interface (ALI) cultures were infected as previously described^12,13^.

### 2.3 5342191 inhibition of virus replication

Compound was dissolved to 10 mM or 1 mM stock concentration in dimethyl sulfoxide (DMSO) and stored at −20°C for subsequent experiments. Compound purity was determined to be >95% as assessed by high resolution mass spectrometry. To examine the effect of 5342191on adenovirus replication, A549 cells were seeded in a 6-well plate at a density of 5 × 10^5^ cells/well. Cells were infected one day post-seeding at an input multiplicity of infection (MOI) of 100 IU/cell of human adenovirus serotype 5 (HAdV-C5). After 1 h of adsorption at 37°C, inoculum was removed and replaced with fresh culture medium containing 1% DMSO (solvent control) or compound dissolved in DMSO (duplicate wells per condition). Progeny virus was harvested 24 h p.i. by scraping the cells into the culture fluid, followed by freeze-thaw of suspension five times with vortexing. The lysate was clarified by centrifugation at 500 x g for 5 min and titrated by endpoint dilution in HEK 293 cells ^15^.

To examine the dose-dependent effects of 5342191 on coronavirus replication, Huh7 cells were seeded in a 96 well plate and infected at an input MOI of 0.1 or 1 TCID_50_ for HCoV-229E and HCoV-OC43, respectively. Similarly, to investigate the dose-dependent impacts of 5342191 on the replication of influenza A virus PR8, A549 cells were seeded in a 96-well plate and infected at an input MOI of 2 TCID_50_. To study the effects of the compound on viral protein levels and viral RNA in media (sups), cells were seeded in a 6 well plate and infected with HCoV-229E, HCoV-OC43, or SARS-Cov2 ^16^ at an input MOI of 0.03, 0.3, and 1 TCID_50_, respectively. For influenza, A549 cells were seeded in a 6-well plate and infected with influenza A virus PR8 at an MOI of 2 TCID_50_. Infection with 229E, OC43, SARS-Cov2, and influenza A virus PR8 inocula was done in serum-free DMEM for an hour with gentle rocking of the plate every 10 min during incubation with virus. Virus inoculum was subsequently removed after an hour, cells washed with 1X PBS, and treated with DMSO or compound diluted in DMEM. For HCoV-229E, HCoV-OC43, and SARS-Cov2 infections, DMEM was supplemented with 2% FBS and 1% penicillin/streptomycin, whereas for PR8 influenza A infection, the media was not supplemented with FBS but with 2.5 μg/mL TPCK trypsin and 1% penicillin/streptomycin.. Cell media were harvested at 1 dpi for PR8, 1 or 2 dpi for HCoV-229E or SARS-Cov2 infection and 4 dpi. for HCoV-OC43 infection, followed by heat inactivation at 95°C for 5 min. For time of addition studies, cells were infected at an input MOI of 2 TCID_50_, DMSO or 5342191 added at various times post virus removal, and media harvested at 24 h post infection to assay viral RNA levels. Media was directly used to detect viral RNA using Luna^®^ Universal One-Step RT-qPCR Kit (NEB) as per manufacturer’s instructions. RT-qPCR assays were set up as follows: 5 µL Luna Universal One-Step Reaction Mix (2X), 0.5 µL Luna WarmStart^®^ RT Enzyme (20X), 0.2 µL of each 5’ and 3’ primers (10 µM), and 1 µL of media (template RNA) in a total reaction volume of 10 µL. Sequences used to detect coronavirus and influenza RNA in media are provided in Supplementary Table S1. The qPCR cycling conditions were as follows: reverse transcription at 55°C for 10 min and initial denaturation at 95°C for 1 min followed by 40 cycles of denaturation at 95°C for 10 s and extension at 60°C for 30 s. The melting curve protocol followed with 15 s at 95°C and then 15 s each at 0.2°C increments between 60°C and 95°C. Melting and standard curves were generated by the CFX Maestro Software (version 1.1, Bio-Rad).

### 2.4 Effect on viral and host protein expression

To assess the effects on viral proteins by western blot, cell extracts were prepared in RIPA buffer (50 mM Tris-HCl pH 7.5, 150 mM NaCl, 1% NP-40, 0.5% sodium deoxycholate, 0.1% SDS) and fractionated on 10% TGX acrylamide stain-free (Bio-Rad) or 14% SDS-PAGE gels. Stain free gels were directly imaged on ChemiDoc MP Imager (Bio-Rad) to measure total protein levels which served as loading control. Following imaging for total protein levels, protein was transferred to PVDF using the Trans-blot Turbo Transfer System (Bio-Rad). Blots were blocked in either 5% Milk-TBS-T (5% Milk, 0.05% Tween-20, 1x TBS) or 3% BSA-TBS-T (3% BSA, 0.05% Tween-20, 1x TBS) prior to incubating with primary antibodies (all diluted in either 5% BSA-TBS-T or 5% TBS-T). Primary antibodies used to evaluate the effect of 5342191 on SR protein and SR kinase expression levels, were; mouse α-SRSF1 (Life Technologies, Cat: 32-4500), rabbit α-SRSF2 (BD Pharmingen, Cat: 556363), mouse α-SRSF3 (Life Technologies, Cat: 33-4200), rabbit α-SRSF4 (Novus Biologicals, Cat: NBP2-04144), rabbit α-SRSF5 (MBL, RN082PW), rabbit α-SRSF6 (Novus Biologicals, Cat: NBP2-04142), rabbit α-SRSF7 (Abcam, Cat: ab137247), rabbit α-SRSF9 (MBL, Cat: RN081PW), mouse α-SRSF10 (Novus Biologicals, Cat: H00010772-M07), rabbit α-Tra2β (Abcam, Cat: ab31353), mouse α-CLK1 (Santa Cruz, Cat: 515897), rabbit α-CLK2 (Abcam, Cat: ab65082), mouse α-CLK3 (Abnova, Cat: H0001198-M05), and rabbit α-SRPK1 (Cedarlane, Cat: OAAN01583). Adenovirus E1A and hexon proteins were detected using rabbit α-E1A (Santa Cruz sc-430) and undiluted hybridoma (2Hx-2) media, respectively, as previously detailed ^17^. To measure the effect of compound on HCoV-229E, HCoV-OC43, or SARS CoV-2 nucleocapsid (N) protein expression, mouse α-coronavirus antibody clone FIPV3-70 (Novus Biologicals, Cat: NB10064754), mouse α-coronavirus group antigen antibody clone 542-7D (Millipore Sigma, Cat: MAB9013), or rabbit anti-SARS CoV2 nucleocapsid antibody (SinoBiological, Cat: 40143-R019) were used, respectively. Changes in HCoV-OC43 spike protein expression were assessed using mouse monoclonal antibody 541-8F (Sigma-Aldrich, MAB9012). The effect of the compound on influenza A virus NS1 non-structural or nucleoprotein expression was measured using rabbit anti-influenza A virus NS1 nonstructural protein antibody (GeneTex, Cat: GTX125990), and rabbit anti-influenza A virus nucleoprotein antibody (GeneTex, Cat: GTX125989), respectively. Following overnight incubation with primary antibodies at 4°C, blots were washed and incubated in appropriate secondary antibody (either α-mouse or α-rabbit horseradish peroxidase (HRP)-conjugated IgG, Jackson ImmunoResearch) for an hour at room temperature. After subsequent washes, signals were detected by ECL (Perkin-Elmer), ECL Plus (Perkin-Elmer), or Clarity Western ECL reagent (BioRad) and imaged using a ChemiDoc MP Imager (Bio-Rad). To test whether changes in protein migration were attributable to altered phosphorylation, extracts were treated with lambda phosphatase (New England Biolabs, Cat: P0753S) prior to running on gels. Relative intensity of detected bands was normalized to corresponding bands of the loading control (GAPDH) or total protein using Bio-Rad ImageLab software. Immunofluorescence and fluorescent *in situ* hybridization analysis of 5342191’s effect on HCoV-229E RNA and protein expression were performed as previously described ^12,13^.

### 2.5 Toxicity assays

Cytotoxicity of 5342191 was assessed using alamarBlue (Life Technologies) or trypan blue exclusion (Life Technologies) and expressed relative to cells treated with DMSO (1%) alone. alamarBlue was added to culture medium prior to harvest/fixation, cells incubated at 37°C in a 5% CO_2_ humidified incubator for 2-6 h, and fluorescence reflecting cell metabolic rate measured using Bio Tek Cytation 5 or TECAN infinite 200Pro fluorescence plate reader (fluorescence detection, 560 nm (ex)/590 nm (em)).

### 2.6 RNA/DNA Analysis

Total RNA was extracted from cells using the BioRad Aurum Total RNA Lysis Kit (BioRad) as per manufacturer’s instructions. Purified RNA (0.5-2 μg) was reverse transcribed using M-MLV (Invitrogen) to generate complementary DNA (cDNA). Viral and β-actin mRNA levels in DMSO- or compound-treated samples were quantified by qPCR as described ^18^. Target quantification was evaluated using the absolute quantification method, normalized to β-actin expression, and expressed relative to DMSO treatment. cDNA was produced from extracted RNA by reverse transcription (RT) with the Bio-Rad iScript cDNA Synthesis Kit, according to the manufacturer’s protocol, using 1µg of RNA in a reaction volume of 20 µL. The conditions were as follows: 5 min at 25℃, 20 min at 46℃ and 1 min at 95℃. Subsequent qPCR analysis of the cDNA, diluted 1/100, was done in technical triplicates using Bio-Rad SsoAdvanced Universal SYBR Green Supermix with primers (Supplementary Table S1) at 500nM final concentration, according to the manufacturer’s protocol. Cycling parameters were 98℃ for 3 min then 40 cycles of 95℃ for 15 seconds, and 60℃ for 30 seconds. Abundance of adenovirus RNA in treated samples, relative to TBP RNA (control), was normalized to RNA abundance at 24h p.i. in 1% DMSO.

For adenovirus DNA analysis, cells were seeded on 6 well plates at a density of 5 x10^5^ cells per well. The next day, cells were infected with HAdV-C5 (input MOI 100 IU/cell) for 1h, inoculum was removed, and replaced with culture medium containing DMSO or 5342191. At the indicated times p.i., culture fluid was removed, and cells were lysed *in situ* with lysis buffer supplied with Norgen Biotek DNA extraction kit 53100. Adenovirus DNA levels were determined using the E2B primer pair (see Supplementary Table S1). Experiments were done using biological duplicates, each measured in technical duplicates. Reactions contained 6 μL of Bio-Rad SsoAdvanced Universal SYBR Green Supermix, 0.5 μL reverse primer (16.5μM), 0.5 μL forward primer (16.5 μM), and 5 μL of DNA sample diluted 1/10 in sterile distilled water. Cycling parameters were: 95℃ for 30 seconds, and 40 cycles of 95℃ for 15 seconds and 60℃ for 30 seconds (as detailed in Bio-Rad SsoAdvanced Universal SYBR Green Supermix user manual). Data was analysed using Bio-Rad Maestro software. To assess effects of compound treatment on viral RNA expression/splicing, assays were performed as previously described ^11^.

### 2.7 Statistical analysis and Visualization

Data is representative of at least three independent experiments, each performed in duplicate, and values presented relative to untreated samples. Statistical differences were assessed by one-way or two-way ANOVA, graphed using Graph Pad Prism 6.0, and results were reported as means ± SDE. P-values < 0.05 were considered statistically significant. For the graphical presentation of the experimental workflow, icons and visual elements were designed using BioRender (www.biorender.com) icon library to enhance clarity and comprehension.

## 3. Results

### 3.1 5342191 inhibits adenovirus replication

As an initial test of 5342191’s ability to affect the replication of another virus dependent on RNA processing for its gene expression and replication, we carried out experiments with adenovirus HAdV-C5. As shown in Fig. 1A, treatment of infected A549 cells with increasing doses of 5342191 resulted in a marked reduction in virus yield, achieving a maximal effect (∼1000 fold reduction in virus yield) at 5 µM, with limited effect on cell metabolism up to 10 µM. Subsequent tests determined that treatment with 5342191 reduced viral DNA accumulation (Fig. 1B) and late viral gene expression (Fig. 1C), consistent with the compound acting at the early stages of viral gene expression, independent of viral DNA replication. Western blot analysis confirmed that 5342191 had a limited effect on the earliest viral protein expressed (E1A) but significantly reduced the expression of a late protein (hexon) (Fig. 1D). These observations suggested that 5342191 does not affect the delivery of viral DNA to the nucleus but rather interferes with subsequent steps in the viral expression cascade necessary for viral DNA replication.

**Figure 1.**
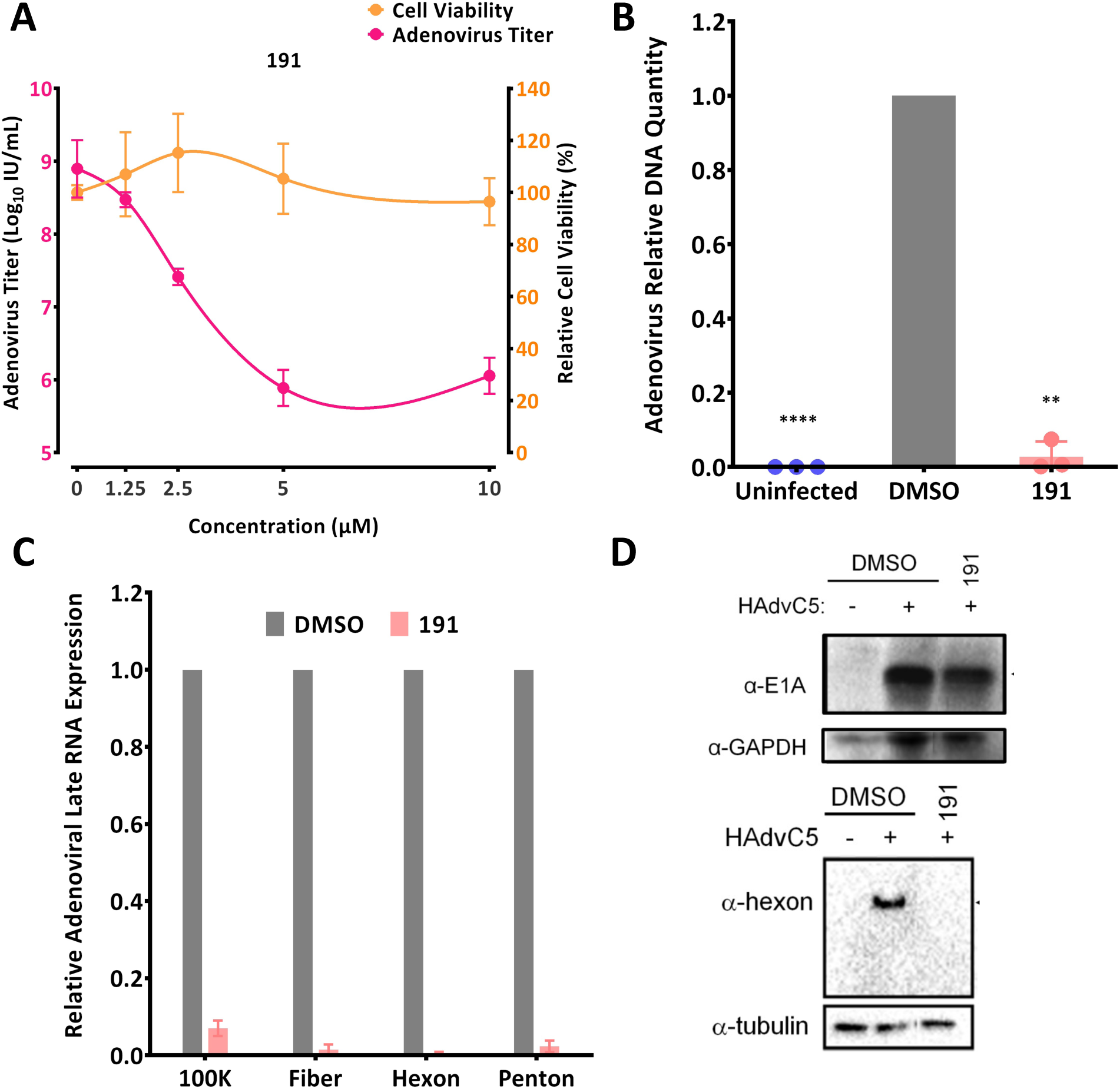
5342191 inhibits adenovirus replication/gene expression. A549 cells were infected with HAdV-C5 at 100 infectious units/cell for 1 h at 37C then virus inoculum was removed, cells were washed with PBS, and fed with fresh medium containing DMSO or 5342191 (191). (A) Samples were incubated with indicated concentrations of 5342191 for 24 hpi, then cells were scaped into the culture fluid and virus was released by freezing and thawing the cell suspension. The cell lysate was clarified by centrifugation and virus yield was measured by endpoint dilution assay. Parallel analysis assessed impact of 5342191 treatment on cell viability using trypan blue or alamarBlue assay. (B) To assess effect of 5342191 on viral DNA amplification, total cellular DNA was harvested 24 hpi from cells treated with either DMSO or 5 µM 5342191 and analysed by qPCR. C) To assess effect of 5342191 on viral late gene expression, total cellular RNA was harvested 24 hpi from cells treated with either DMSO or 5 µM 5342191 for detection of selected adenovirus transcripts by RT-qPCR. (D) To assess effect of 5342191 on viral protein expression, cell lysates were prepared 24 hpi from cells treated with either DMSO or 5 µM 5342191 and levels of adenoviral E1A and hexon determined by immunodetection on a western blot. Data shown in this figure represents results from n=3 assays.

To identify the specific stage of viral gene expression affected, we examined the impact of 5342191 on several early viral genes (E1A, E1B, E2A, and E2B) by RT-qPCR. Consistent with its limited effect on E1A protein levels, 5342191 had limited impact on E1A RNA accumulation at doses sufficient to suppress virus replication (Fig. 2A). In contrast, 5342191 caused a dose dependent reduction in accumulation of E1B, E2A, and E2B RNAs (Fig. 2A). To gain further insight, we also examined for changes in E1A RNA alternative splicing. As observed previously with both GPS491 and cardiotonic steroids ^11,17^, 5342191 addition blocked the time dependent shift in E1A RNA splicing from predominantly 13S/12S early after infection to predominantly 9S at later time (16, 24 hpi) (Fig. 2B) in a dose-dependent manner.

**Figure 2.**
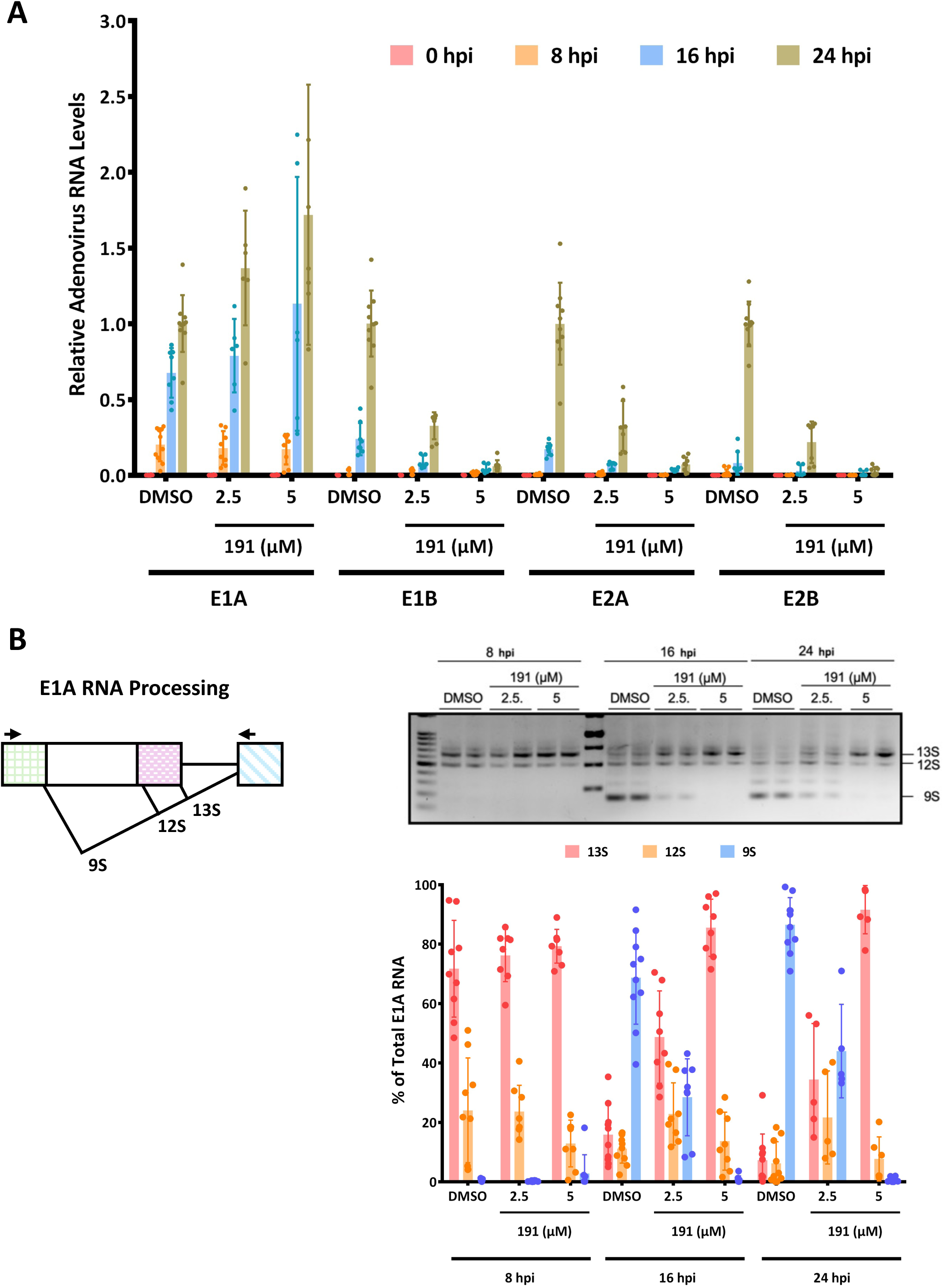
5342191 affects adenovirus RNA accumulation/splicing. A549 cells were infected with HAdV-C5 at 100 infectious units/cell for 1 h at 37C then virus inoculum was removed, cells were washed with PBS and fed with fresh medium containing DMSO or 5342191 (2.5 or 5 uM). Total RNA was harvested at 8 hpi, 16 hpi, and 24 hpi then assessed for changes in (A) viral early (E1A, E1B, E2A, E2B) RNA accumulation by RT-qPCR or (B) alterations in adenoviral E1A RNA splicing by RT-PCR. At left, a schematic of E1A RNA splicing, at right a representative gel of patterns of E1A RNA isoforms (13S, 12S, 9S) observed upon treatment with DMSO or 5342191 (2.5 or 5 µM) and at bottom, results pooled from n=3 assays.

### 3.2 53421 inhibits replication of multiple human coronaviruses

The capacity of 5342191 to inhibit adenovirus replication led us to question whether, similar to GPS491 and cardiotonic steroids ^11,13^, it could also be effective against coronaviruses. To test this hypothesis, we examined the dose-dependent effect of 5342191 on the replication of HCoV-229E in Huh7 cells. As shown in Fig. 3B, 5342191 is a potent inhibitor of HCoV-229E replication with an EC_50_ of 1.25 µM with limited toxicity in this system at 12.5 µM. Subsequent tests confirmed that the reduction in viral RNA accumulation in the media seen in Fig. 3B upon 5342191 addition was accompanied by a significant reduction in infectious virus release; a dose of 5342191 sufficient to reduce HCoV-229E RNA accumulation in media by 50% resulted in over 90% reduction in infectious virus yield (Fig. 3C). Similar observations were made upon HCoV-OC43 coronavirus infection, where 2.5 μM of 5342191 led to over 90% inhibition of viral release into the media (Fig. 3D). Further tests determined that the addition of 5342191 1 hour post-infection (hpi) not only blocked viral RNA release in the media 24 hpi (Fig. 3D) but also inhibited intracellular accumulation of both genomic and total viral RNAs (Fig. 3E) and viral proteins (Fig. 3F).

**Figure 3.**
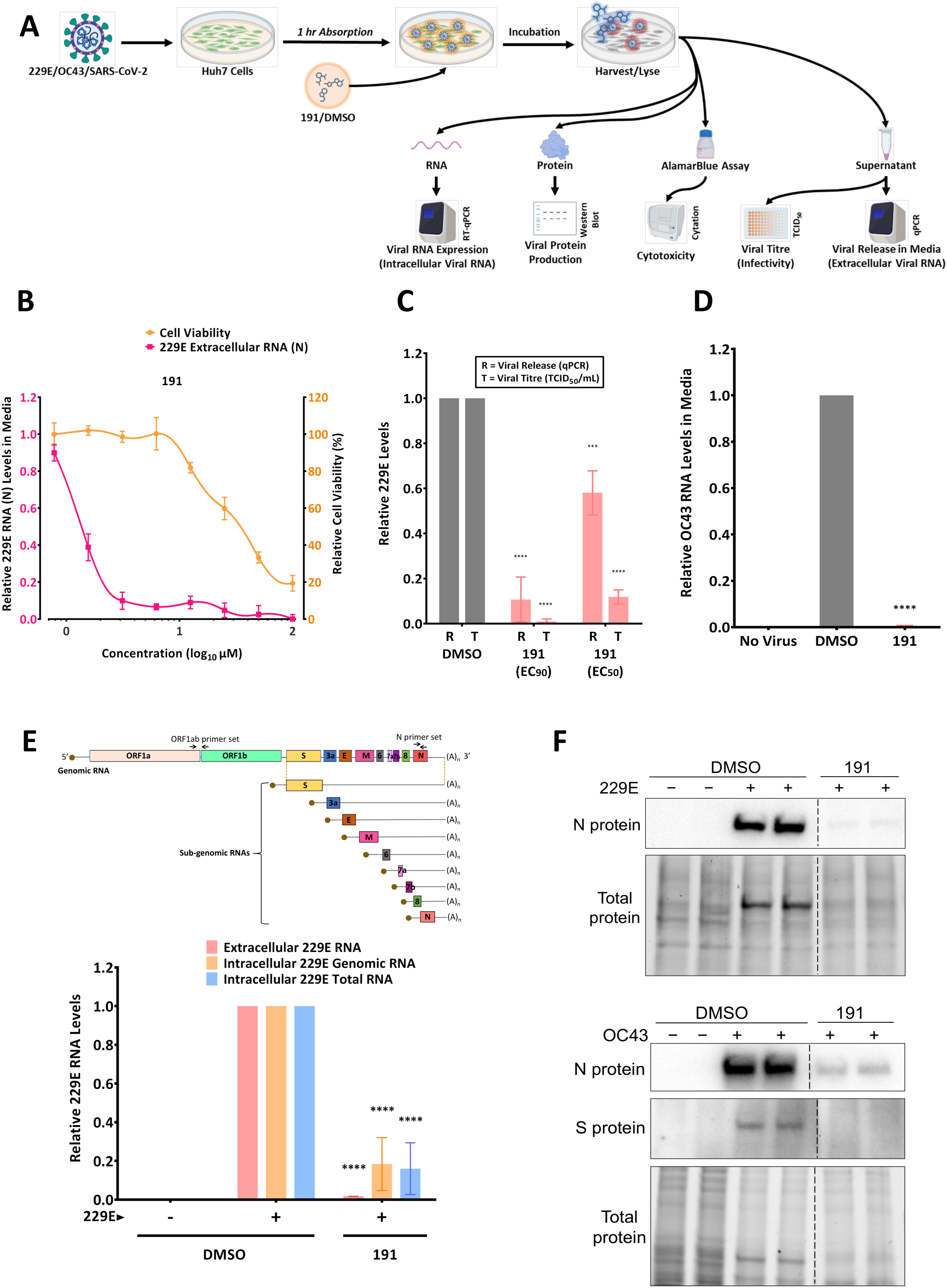
5342191 inhibits HCoV-229E and HCoV-OC43 replication. (A) Schematic of experimental protocol. Huh7 cells were infected with indicated coronavirus strains for 1 h, cells washed, then treated with either DMSO or 5342191. Media was harvested and levels of viral RNA release into media determined by RTqPCR. Cells were lysed for RNA extraction to measure levels of intracellular viral RNA using RT-qPCR, and viral protein production via western blotting. (B) Effect of 5342191 concentrations on HCoV-229E RNA accumulation in media (by RTqPCR) and cell viability (assessed using Alamar blue). (C) Correlation of viral RNA release and infectious virus produced. Infected cells were treated with EC_50_ (1.2 µM) or EC_90_ (2.5 µM) doses of 5342191 for 24 h then media harvested and viral RNA in media accumulation measured by RT-qPCR, infectious virus by TCID_50_ assay. (D) Cells were infected with HCoV-OC43 virus at an MOI of 2 and then treated with either DMSO or 2.5 µM 5342191 for 24 hours. Culture supernatants were harvested and analyzed by RT-qPCR to measure the release of HCoV OC43 into the culture media. (E) Top, schematic of coronavirus virus genome and subgenomic RNAs generated. Shown are the position of primer sets used to measure genomic or total viral RNAs. Bottom, infected cells were treated with DMSO or 2.5 µM 5342191 for 24 h. Extracellular viral RNA accumulation was measured by RT-qPCR. Total RNA was extracted from cells and abundance of genomic or total viral RNA (genomic and nested RNAs) measured by RT-qPCR. (F) Representative immunoblots showing the impact of 5342191 on the production of the nucleocapsid (N) protein of the HCoV-229E virus (top) and the N protein and Spike (S) protein of the HCoV-OC43 virus (bottom).

Due to the similar effects observed with HCoV-229E and HCoV-OC43, we investigated whether 5342191 was also effective against SARS-CoV-2. Using a 10-fold serial dilution of compound, we found that even at concentrations as low as 300 nM, 5342191 significantly suppressed the release of SARS-CoV-2 by approximately 80%. (Fig. 4A). Tests in Huh7 cells confirmed that 534219 effectively suppressed the replication of multiple SARS-CoV-2 strains (SB2, Delta, Omicron) as assessed by viral RNA release into the media (Fig. 4B) and intracellular viral RNA accumulation (Fig. 4C). 534219 addition also reduced SARS-CoV-2 nucleocapsid (N) protein expression by approximately 90% (Fig. 4D). As a further test of 5342191’s inhibitory activity, we also examined its effect on HCoV-OC43 and SARS-CoV-2 replication in human lung organoids (Fig. 5A). As shown in Fig. 5B-C, both HCoV-OC43 and SARS-CoV-2 can infect this tissue, and their replication was effectively blocked upon the addition of 5342191 addition post-infection.

**Figure 4.**
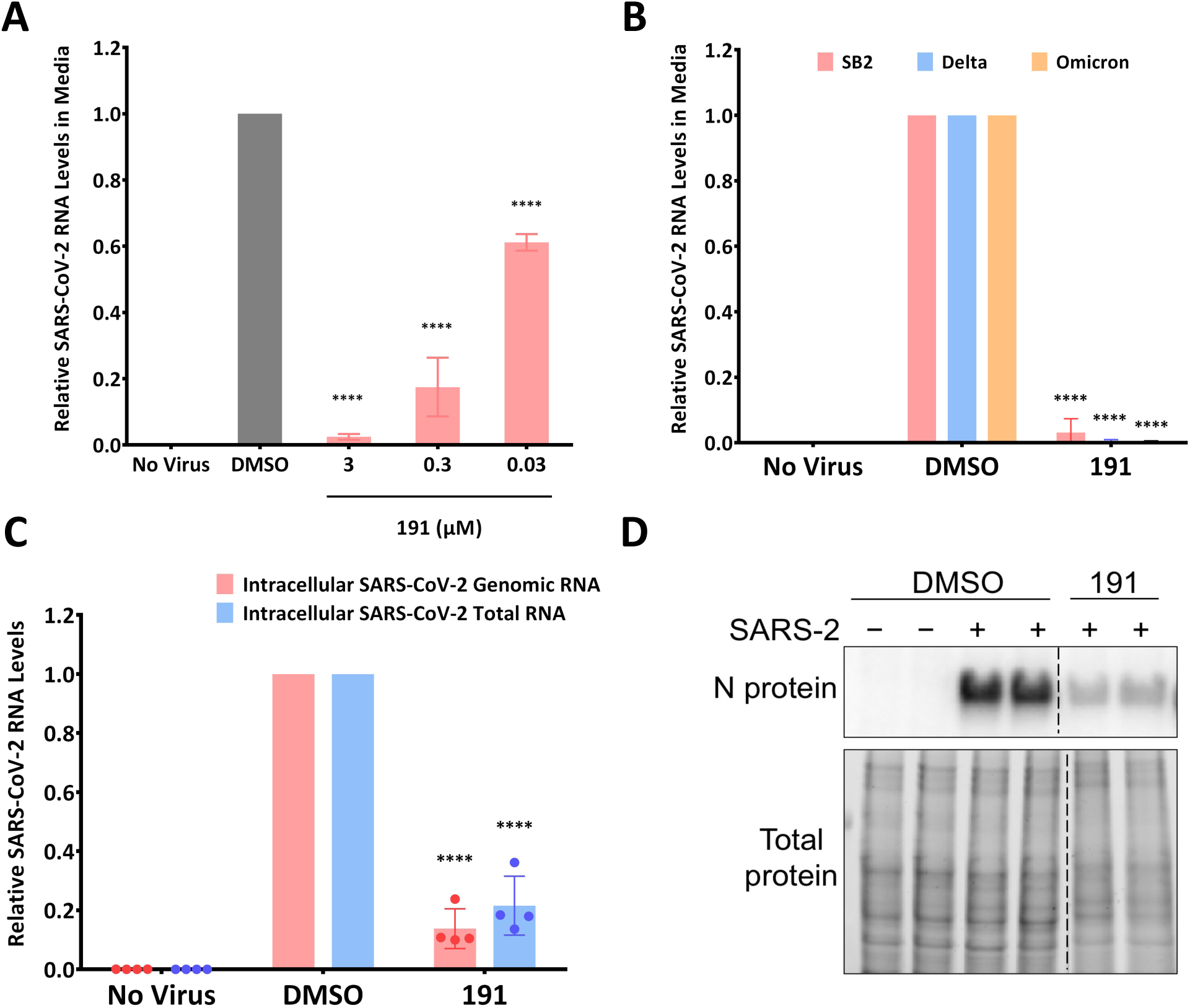
5342191 inhibits replication of SARS-CoV2 strains SB2, Delta, and Omicron. (A) Effect of 10-fold dilutions of 5342191, starting from 3 µM to 30 nM, on the release of SARS-CoV-2 SB2. Supernatants were harvested 24 hours after infection of Huh7 cells and analyzed by RT-qPCR. (B). Huh7 cells were infected with SARS-CoV2 SB2, Delta, or Omicron (MOI=2) for 1 h, cells washed, then treated with either DMSO or 2.5 µM 5342191. Media was harvested at 24 hpi and RTqPCR used to measure viral release. (C) RNA samples were extracted from SARS-CoV-2-SB2-infected Huh7 cells and measured by RT-qPCR to determine levels of total (N) and genomic viral RNA. (D) Representative immunoblot showing the impact of 5342191 on the expression of the nucleocapsid (N) protein of SARS-CoV-2 SB2. Results shown are the average of n=3 independent assays.

**Figure 5.**
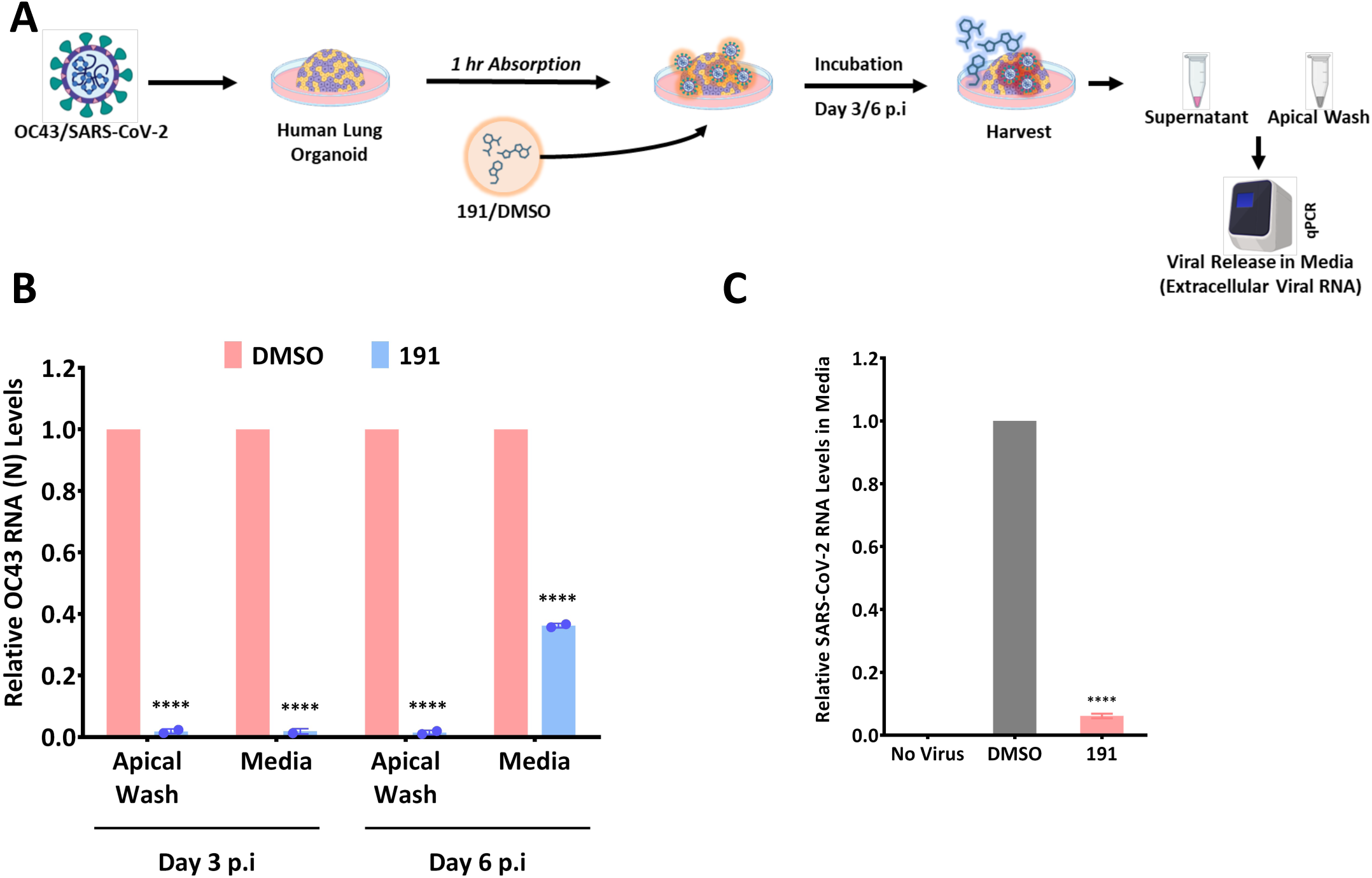
5342191 inhibits replication of HCoV-OC43 and SARS-CoV2 in human lung organoids. (A) Outline of experiment. Human lung organoids were infected with HCoV-OC43 virus (B) or SARS-CoV2 SB2 at a MOI of 2 for 1 h. Organoids were then washed several times to remove virus inoculum then incubated with media containing DMSO or 2.5 µM 5342191. Three or six days post-infection, media was harvested and level of viral RNA determined by RTqPCR. Shown are the results from n>3 independent assays.

### 3.3 5342191 acts post virus entry to preferentially inhibit nested HCoV-229E RNA production

Having established that 5342191 inhibits coronavirus replication, we investigated the basis for the response. Our previous studies demonstrated that, in Huh7 cells infected with HCoV-229E at a multiplicity of infection (MOI) of 2, both intracellular viral RNA and protein were first detectable at 9 hpi with viral replication foci becoming visible by 16 hpi^11–13^. To determine whether 5342191 acts post virus entry, we examined the effect of delayed compound addition on virus replication. As shown in Fig. 6A, 5342191 addition at 12 or 16 hpi was almost as effective at inhibiting virus release as addition at 1 hpi. This suggests that the compound targets a post-entry step, even after the viral replication complex has formed.

**Figure 6.**
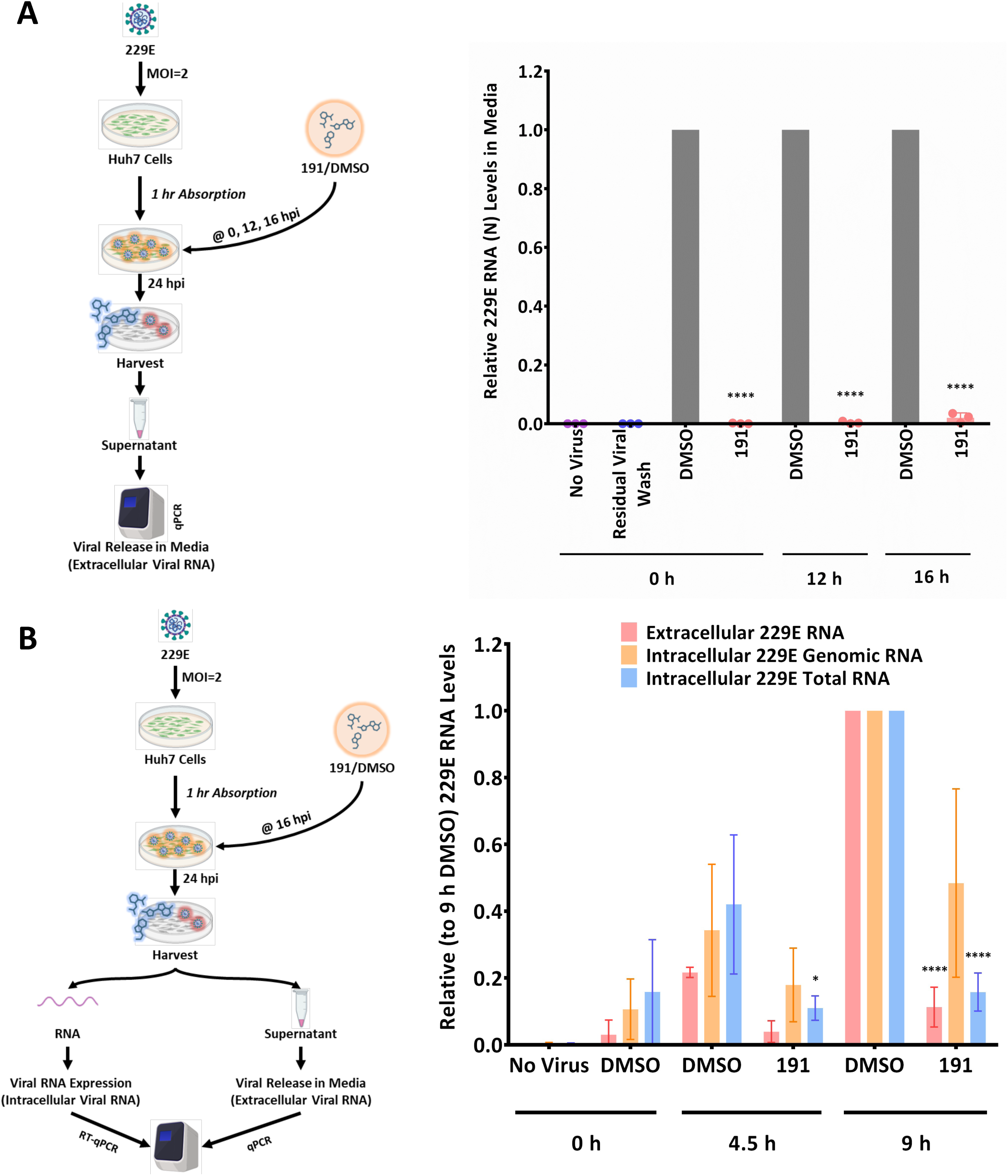
5342191 acts post virus entry to preferentially inhibit HCoV-229E nested RNA synthesis. (A) Huh7 cells were infected with HCoV-229E (MOI 2) for 1 h then media replaced. 5342191/DMSO was added 0, 12hpi or 16 hpi and incubation continued. At 24 hpi, media was harvested, and viral levels determined by RTqPCR. (B) Huh7 cells were infected with HCoV-229E (MOI 2) for 1 h then media replaced. At 16 hpi, media was replaced with media containing DMSO or 5342191 and incubation continued for, 0, 4.5, or 9 h at which time media and cells were harvested and indicated viral RNA levels determined by RT-qPCR. Total viral RNA = genomic + nested viral RNAs. Shown are the results from n>3 independent assays.

To further explore how 5342191 impacts virus replication, we assessed how the compound’s delayed addition affects viral RNA accumulation. Coronavirus replication involves the amplification of genomic RNA as well as the generation of a series of nested RNAs that encode the viral structural proteins^19–22^. To assess whether 5342191 affects the synthesis of all viral RNAs equally, we examined the effect of delayed compound addition on the accumulation of both genomic and total (a sum of genomic and nested) viral RNAs. As indicated in Fig. 6B, cells were infected at an MOI of 2 for 1h, the virus inoculum was removed, and cells were incubated for 16 h before adding DMSO or 5342191. Total RNA was extracted 0, 4.5, or 9 h post compound addition, and the abundance of viral RNAs determined. While the accumulation of viral genomic RNA was reduced by ∼50 % at 9 h after 5342191 addition, there was limited or no increase in total viral RNA abundance over the same period. Given that the primer set used in the RT-qPCR for total viral RNA is in the N region common to all viral RNAs, the signal obtained reflects both genomic and nested RNAs. The observed increase in abundance of genomic but not total viral RNA upon 5342191 addition indicates that the synthesis of nested RNAs is preferentially affected.

To further explore the effect of 5342191 on viral RNA synthesis, we also examined its effect on viral replication complex stability. As outlined in Fig. 6, Huh7 cells were infected with HCoV-229E (MOI 2), the virus inoculum removed, and cells incubated for 16 h before adding DMSO or 5342191. Cells were then fixed 0 or 9 h post-compound addition and processed for immunofluorescence or in situ hybridization. Viral replication centers are detected due to the accumulation of dsRNA (a replication intermediate) within perinuclear invaginations of the endoplasmic reticulum^19,23,24^. Immunofluorescent imaging (Fig. 7A) revealed that the pattern of dsRNA staining was unaffected by 5342191 addition (DMSO 24 h vs. 5342191 24h), indicating that the replication complexes remained stable 9 h after compound addition. Parallel analysis of viral RNA localization by *in situ* hybridization revealed a similar effect (Fig. 7B). While the signal for total viral RNA was dispersed throughout the cytoplasm, genomic RNA remained largely within perinuclear foci regardless of whether cells are treated with DMSO or 5342191.

**Figure 7.**
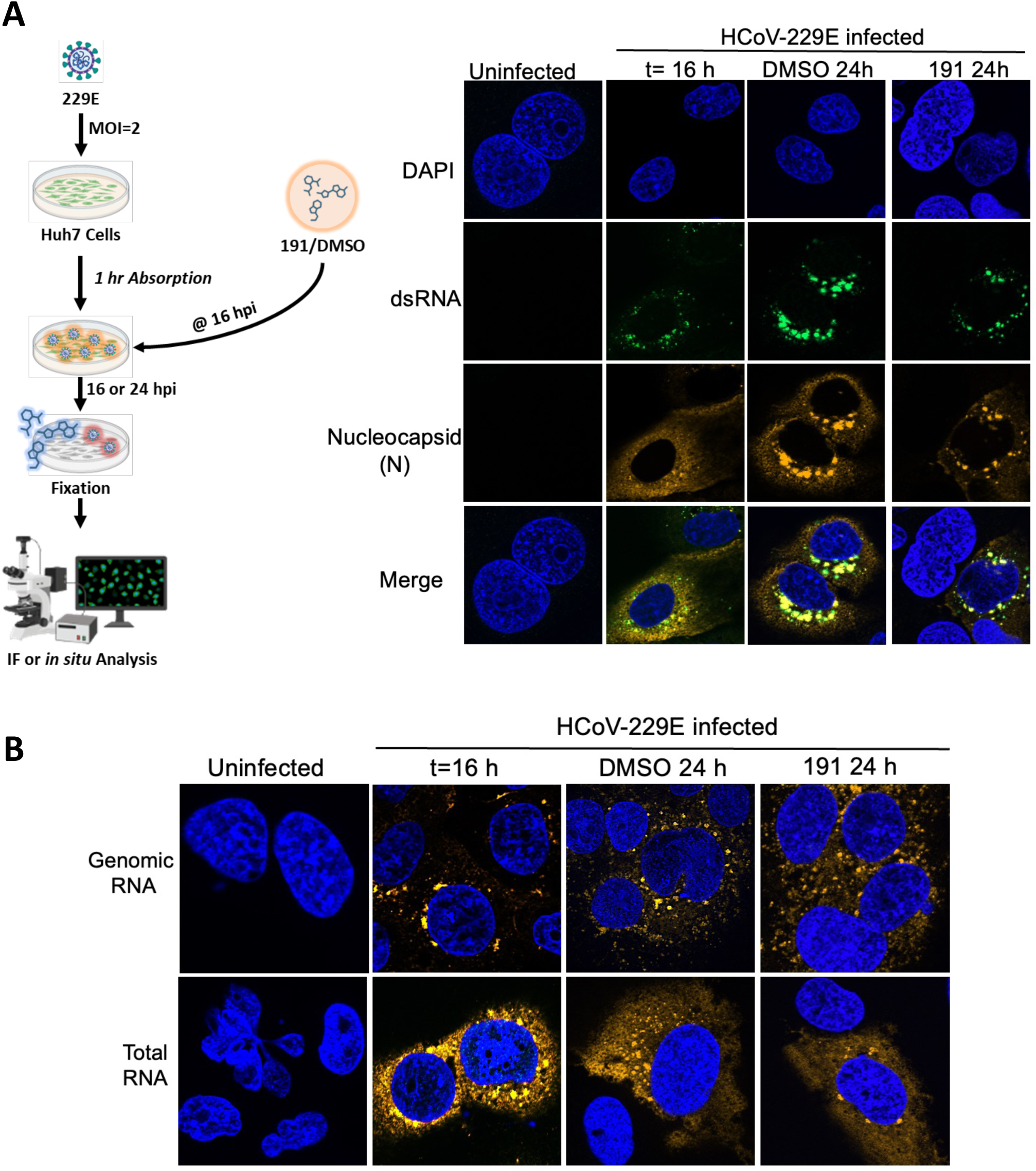
5342191 does not disrupt HCoV-229E replication centers. Huh7 cells were infected with HCoV-229E (MOI 2) for 1 h then media replaced. At 16 hpi (t=16 h), media was replaced with media containing DMSO or 2.5 µM 5342191 and incubation continued for 9 h (DMSO 24h, 191 24h) at which time cells were fixed and (A) stained to detect viral replication centers (dsRNA) or viral nucleocapsid (B) processed for in situ hybridization using probes detecting viral genomic or total RNAs. Shown are representative images from n=3 independent assays.

### 3.4 5342191 inhibits PR8 influenza A virus replication

The ability of 5342191 to inhibit one group of viruses with pandemic potential led us to explore its effectiveness against other viruses, such as influenza. To this end, we examined the effects of 5342191 on influenza PR8 replication in A549 cells. Similar to its effects on coronaviruses, 5342191 reduced PR8 virus accumulation in the media (as measured by both TCID_50_ assays or RT-qPCR for viral RNA) in a dose dependent manner with an EC_50_ of 51 nM and a CC_50_ >10 µM (Fig. 8A-B). Subsequent analysis of intracellular viral nucleoprotein (NP) and non-structural protein 1 (NS1) (Fig. 8C), or viral RNA (Fig. 8D), showed a 90% reduction in the accumulation of all measured viral proteins and RNAs. Additionally, accumulation of spliced forms of M and NS RNAs (M2, NS2) was also reduced (Fig. 8E) upon treatment with an EC_90_ dose of the compound. To define which stage of PR8 replication was impacted by 5342191, we assessed the compound’s effectiveness when added several hours after infection. We first established the kinetics of PR8 replication in A549 cells, measuring both intracellular and extracellular viral RNA and protein accumulation. As detailed in Fig. 9A, extracellular viral RNA was first detected at 12 hpi, peaking at 24 hpi at an MOI of 2. Parallel analyses revealed a gradual accumulation of intracellular viral proteins (Fig. 9B) and RNAs (Fig. 9C) with a significant increase in abundance above baseline at 12 hpi. Based on these observations, we explored whether delayed addition of 5342191 affected virus production. Similar to its effects on HCoV-229E, adding 5342191 at 1 or 6 hpi resulted in equivalent reductions in virus yield in the media. Addition at 9 or 12 hpi compromised virus release to some extent (Fig. 9D), consistent with 5342191 affecting a post-entry step of virus replication.

**Figure 8.**
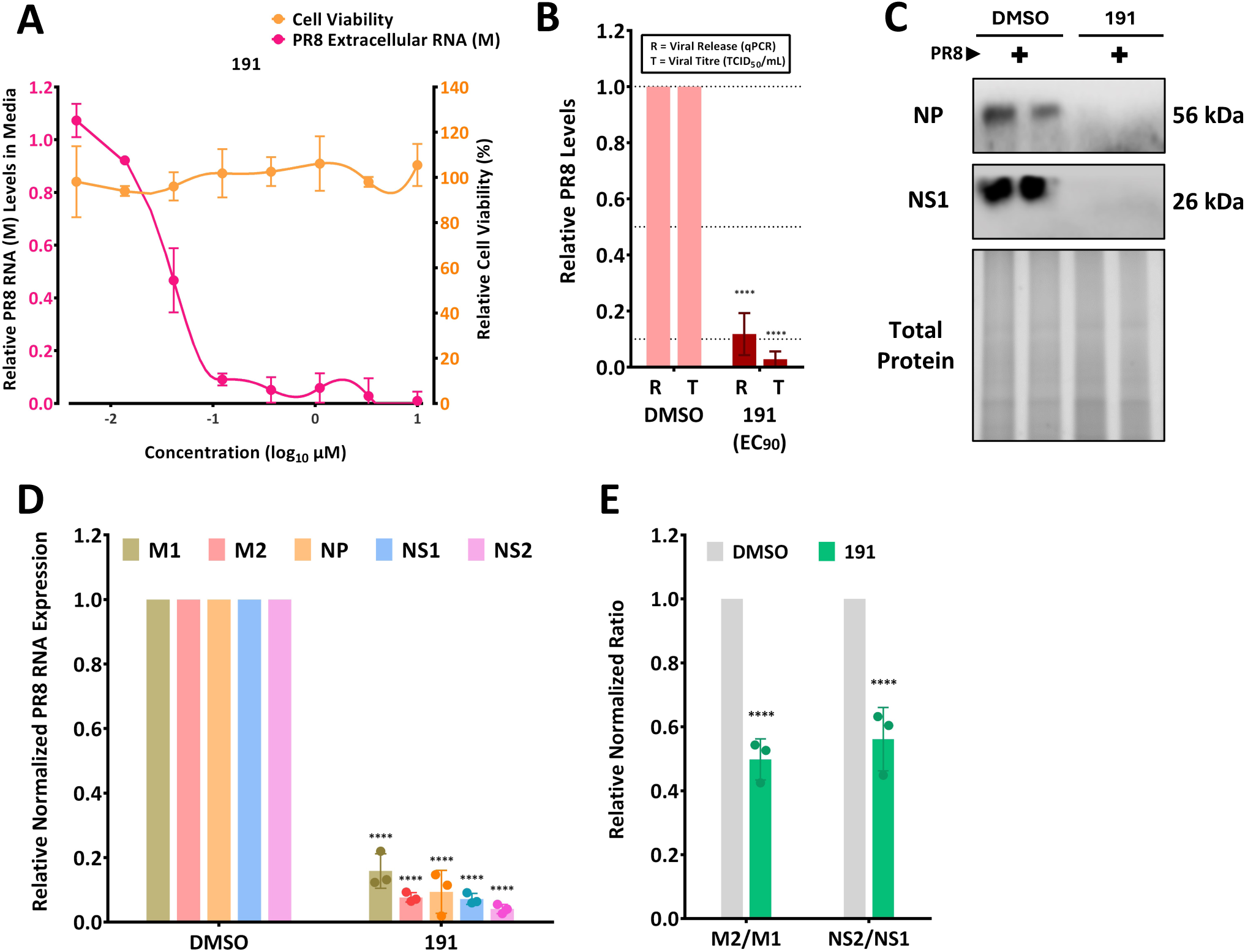
5342191 inhibits PR8 influenza A virus replication. (A) A549 cells were infected at an MOI of 2 with PR8 for 1h, media removed and replaced with media containing DMSO or indicated doses of 5342191. At 24 hpi, media was harvested, and levels of viral RNA released determined by RTqPCR. (B) To correlate the level of viral RNA released to infectious virus, infected cells were treated with doses of 5342191 required to reduce PR8 RNA accumulation in media by 50% (EC_50,_ 0.051 µM) or 90% (EC_90,_ 0.104 µM). Media was assessed in parallel to determine the effect of compound addition on the level of infectious virus present as determined by TCID_50_ assay. (C-E) A549 cells were infected with PR8 at an MOI of 2 for 1h, media removed and replaced with media containing DMSO or 0.104 µM 5342191. 24 hpi, cells were harvested, (C) expression of viral NP or NS1 assessed by western blot or (D) total RNA extracted, and intracellular levels of indicated viral RNAs determined by RT-qPCR. (E) Ratio of M2 versus M1 or NS2 versus NS1 to assess impact of 5342191 on viral RNA splicing. Shown are the results from n>3 independent assays.

**Figure 9.**
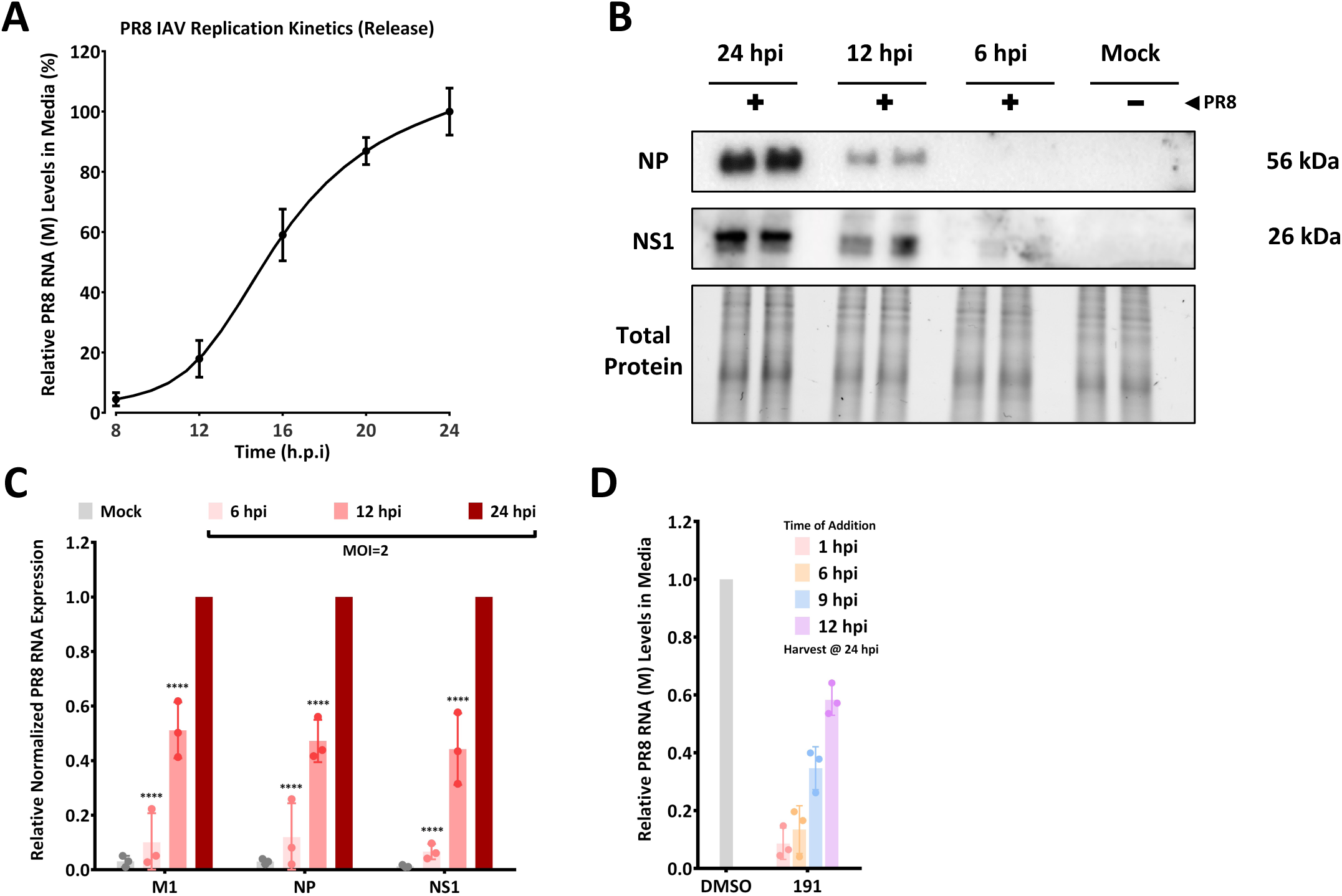
5342191 acts post-entry to inhibit influenza replication. A549 cells were infected at an MOI of 2 with PR8 for 1h, then media replaced. Media was harvested at various times post infection and (A) level of viral RNA in media determined by RTqPCR, (B,C) cells harvested and (B) protein lysates used for western blot to detect PR8 NP or NS1 proteins or (C) total RNA extracted and intracellular abundance of viral RNAs (M1, NP, and NS1) determined by RT-qPCR. (D) To assess whether 5342191 could inhibit virus replication at times after infection, cells were infected with PR8, MOI 2, for 1h, then 0.104 µM 5342191 added 1, 6, 9, or 12 hpi. At 24 hpi, media was harvested, and levels of viral RNA determined by RTqPCR.

To gain further insight into how 5342191 addition affected virus production, we assessed its impact on viral RNA synthesis. As described for HCoV-229E, A549 cells were infected with PR8 (MOI 2) for 1 h, after which the virus inoculum was removed, and fresh media was added. At 9 hpi, DMSO or 5342191 was added, and total RNA was extracted 5.5h post-compound addition. Prior to harvest, cells were treated with ethynyl-uridine (EU) for 1.5 h to label newly synthesized RNA. Click chemistry was used to biotin tag extracted RNAs containing ethynyl-uridine, which were then isolated using streptavidin (Fig. 10A). The abundance of modified and unmodified viral RNAs was determined by RT-qPCR (Fig. 10A) ^25^. Results (Fig. 10B,C) revealed that 5342191 addition significantly reduced accumulation of all viral RNAs examined with the greatest effect on the production of ethynyl-uridine labeled RNAs, indicating inhibition of *de novo* viral RNA production.

**Figure 10.**
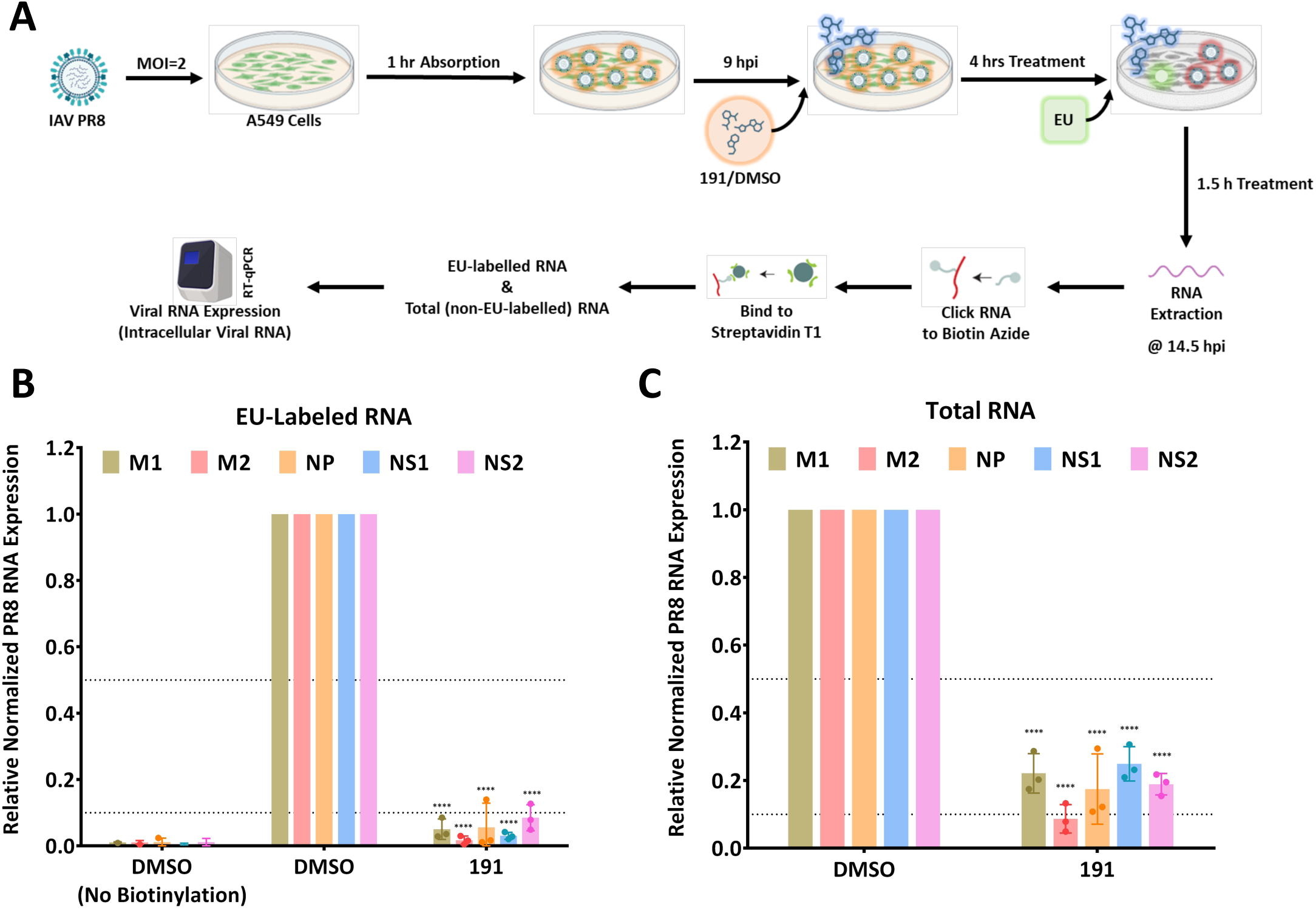
5342191 inhibits influenza RNA synthesis. (A) A549 cells were infected at an MOI of 2 with PR8 for 1h, then media replaced. At 9 hpi, DMSO or 0.104 µM 5342191 was added. 90 minutes prior to harvest, cells were pulse labelled with ethynyl-uridine (EU) to label newly synthesized RNA. RNA was extracted at 14.5 hpi and EU-labelled RNAs biotinylated using click chemistry and isolated using streptavidin beads. Both (B) EU-labelled (EU-RNA) and (C) unlabelled (Total) RNAs were used in RT-qPCR reactions to evaluate abundance of several viral RNAs.

### 3.5 Both HCoV-229E and influenza PR8 have limited capacity to develop resistance to 5342191

One limitation of most DAAs is the rapid emergence of drug-resistant viral variants due to the high mutation frequency of most RNA viruses. To evaluate whether 5342191 could provide more robust control over HCoV-229E and influenza PR8 replication, we tested its ability to select for resistant variants. Briefly (Fig. 11A), cells were infected with HCoV-229E or influenza PR8 at an MOI of 2 for 1 h. The virus inoculum was then replaced with media containing an EC_90_ dose of 5342191 or a DAA (boceprevir or molnupirevir for HCoV-229E, favipiravir or oseltamivir for influenza PR8). Cells were incubated until cytopathic effects were detected, after which media was harvested, diluted ten-fold, and an aliquot was used to inoculate a fresh set of cells for 1 h, followed by addition of EC_90_ dose of the compound. After 5 or 10 rounds of selection, the collected virus was titered (by TCID_50_), and equivalent infectious units were used to infect naïve cells at an MOI of 0.01. One hour post infection, cells were treated with DMSO or EC_90_ dose of drug, and the kinetics of virus release monitored by RT-qPCR. Comparison of the replication kinetics of original (WT) and selected virus in the presence and absence of drug provides a measure of resistance of the selected viruses. As shown in Fig. 11B-C, treatment of WT virus with EC_90_ doses of any of the compounds resulted in a marked reduction in viral RNA accumulation in the media. In contrast, both HCoV-229E and influenza PR8 viruses selected using DAAs after only 4 rounds displayed diminished or no effect of the compound on virus growth kinetics. In contrast, HCoV-229E or influenza PR8 viruses selected with 5342191 remained sensitive to an EC_90_ dose of the compound after ffour rounds of selection (Fig. 11). In the case of HCoV-229E, even 10 rounds of selection did not result in outgrowth of a variant with significant resistance to 5342191 (data not shown). Consequently, compared to the DAAs tested, 5342191 appears to establish a more robust barrier to virus replication as reflected in the greater challenge for the virus to develop resistance.

**Figure 11.**
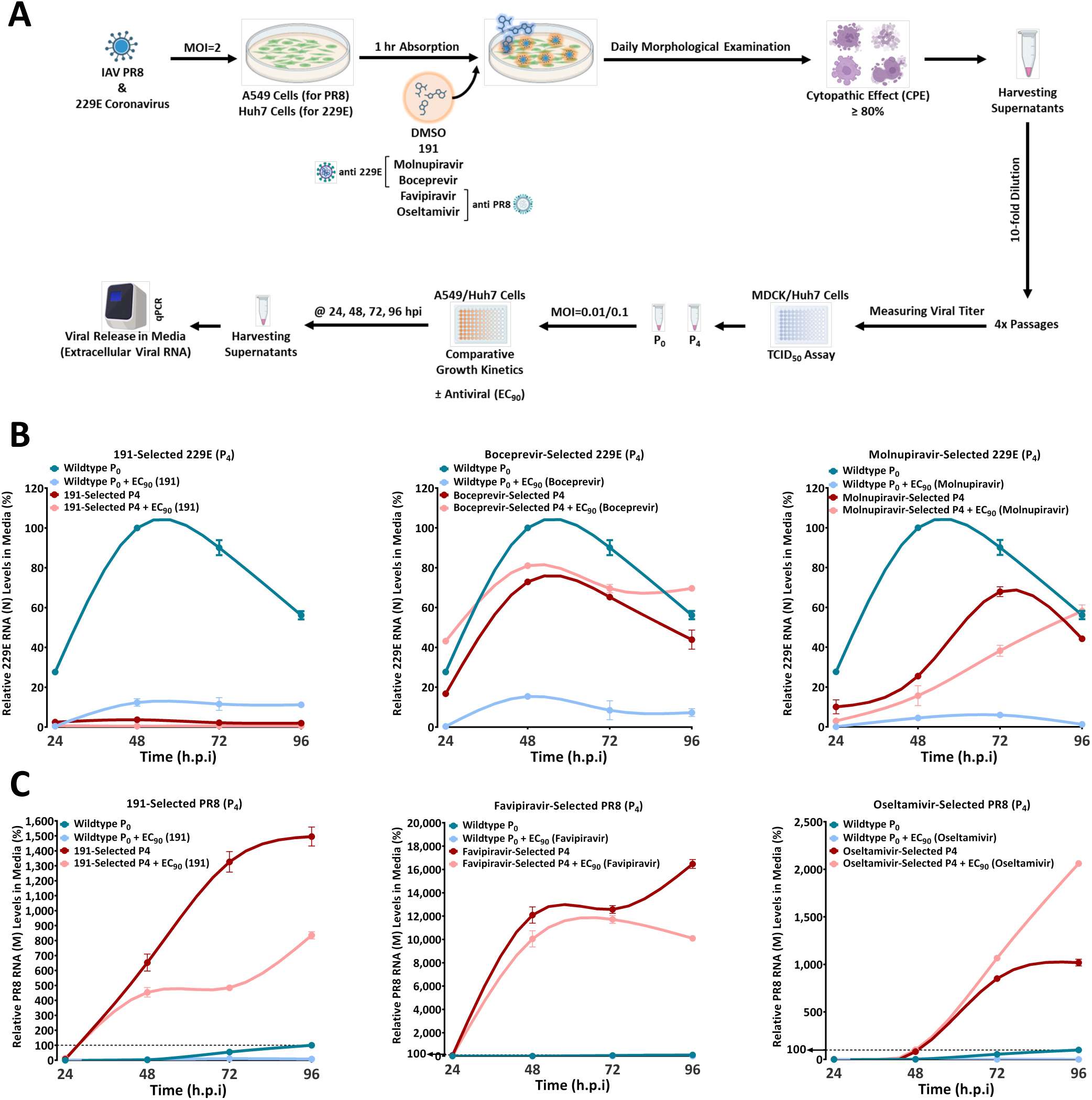
Limited capacity to select for virus resistant to 5342191. (A) Scheme for selection of compound resistant virus. Cells were infected with either (B) HCoV-299E or (C) influenza PR8 then incubated in media containing EC_90_ concentration of compound until cytopathic effect (CPE) was observed. Collected media was diluted ten-fold and used to infect a fresh set of cells. Results shown are after 4 serial passages. After 4 passages, titers of infectious virus in collected media was determined by TCID_50_ assay and equivalent infectious units of original or selected virus used to infect fresh cells at an MOI of 0.01 (PR8) or 0.1 (HCoV-229E). Media was harvested at indicated times after infection and virus replication kinetics assessed by measuring levels of viral RNA release by RTqPCR. Shown are examples of virus growth kinetics for samples run in duplicate (B) HCoV-229E selection with 5342191 (2.5 µM), boceprevir (1.2 µM), or molnupiravir (0.137 µM), (C) influenza PR8 selection with 5342191 (0.104 µM), favipiravir (6.2 µM), or oseltamivir (1.5 µM).

### 3.6 Antiviral effect of 5342191 is blocked by inhibitors of the MAPK pathway or GPCR signaling

Previous investigations into how 5342191 impedes HIV-1 replication revealed a dependence on GPCR/MEK/ERK signaling pathways for its anti-HIV-1 effect^14^. To explore if the anti-coronavirus and anti-influenza activities of 5342191 displayed a similar dependence on these cellular signaling pathways, the ability of various pathway inhibitors to reverse the 5342191 inhibition was tested. Cells were pretreated with DMSO or inhibitors of MEK/ERK, JNK, p38, NCX, or GPCR signaling for 4 h at doses that, on their own, had minimal effect on virus replication (Fig. 12, DMSO). After removing the media, cells were infected with either HCoV-229E or influenza PR8 (MOI 2) for 1 h. The media was then replaced with fresh media containing either DMSO or 5342191 (EC_90_ dose) with or without the previously used pathway inhibitors. At 24 hpi, the media was harvested, and virus RNA release was assessed using RT-qPCR. Results (Fig. 12, 5342191) revealed that, except for the sodium-calcium exchange (NCX) inhibitor, all the tested inhibitors reversed 5342191’s inhibition of HCoV-229E (Fig. 12B) and influenza PR8 (Fig. 12C).

**Figure 12.**
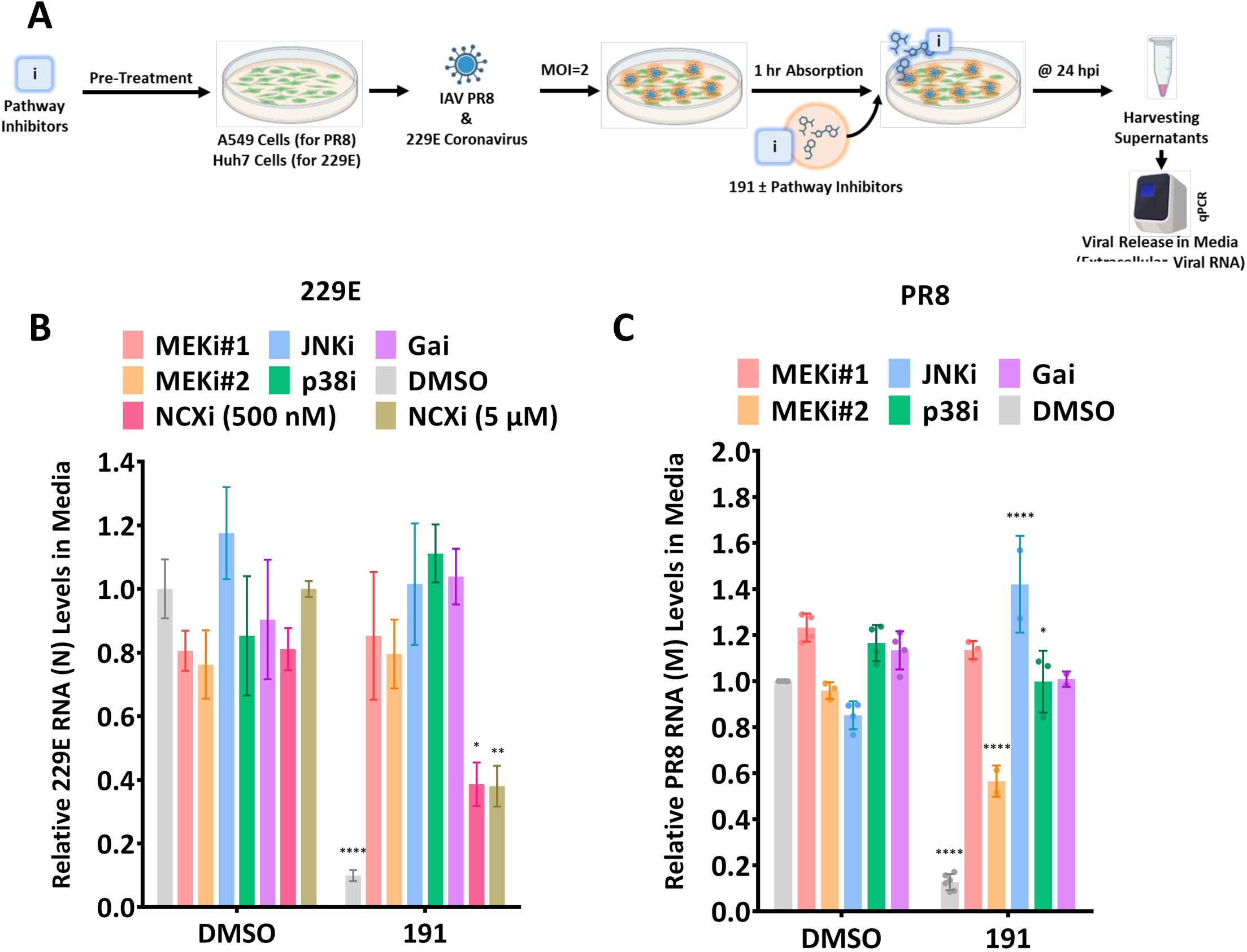
MAPK/GPCR inhibitors reverse antiviral effect of 5342191. (A) Huh7/A549 cells were pretreated for 4 h with indicated pathway inhibitors, washed, infected with virus (MOI 2) for 1 h, then media replaced containing either DMSO or EC_90_ dose of 5342191 (0.104 µM against PR8 and 2.5 µM against 229E) +/− pathway inhibitor. Inhibitors used are as follows; MEKi#1(U0126): 0.250 (PR8) or 0.625 (229E) µM, MEKi#2 (Selumetinib/AZD624): 0.500 (PR8) or 0.625 (229E) µM, Gαi (BIM-46187):0.125 (PR8) or 0.050 (229E) µM, JNKi (SP600125): 0.250 (PR8) or 0.312 (229E) µM, p38i (SB203580): 0.015 (PR8) or 0.450 (229E) µM, NCXi (KB-R7943): 0.5 or 5 µM. 24 hpi, media was harvested, and level of virus release determined by RTqPCR. Summary of results for (B) HCoV-229E or (C) influenza PR8 is shown, n=3 independent assays.

### 3.7 5342191 alters cellular SR protein function and SR kinase expression

Replication of HIV-1, adenovirus, and influenza is critically dependent on proper viral RNA splicing which is altered in response to 5342191 treatment (Figs. 2B, 8E). Changes in HIV-1 RNA accumulation correlate with 5342191-induced changes in the abundance of select SR proteins (reduced SRSF1 and 3, increased SRSF4) known to regulate RNA splicing^14^. To investigate whether 5342191 affects the activity of select SR proteins, we examined its impact on the splicing of a model RNA, Bcl-x. HEK-293 cells were transfected with a vector generating Bcl-x RNA and plasmids expressing individual SR proteins, then treated with DMSO or 5342191. Total RNA was extracted, and levels of spliced isoforms of Bcl-x (Bcl-x*_l_* or Bcl-x*_s_*) were determined by RT-PCR (Fig. 13A). As shown in Fig. 13B-C, while 5342191 had no impact on the level of Bcl-x RNA splicing in EcR293 cells, it significantly affected the ability of SRSF3, 4, 7, 9, 10 and Tra2a to control Bcl-x_s_ production, No significant impact of the compound was detected on SRSF1, SRSF2 and Tra2b.

**Figure 13.**
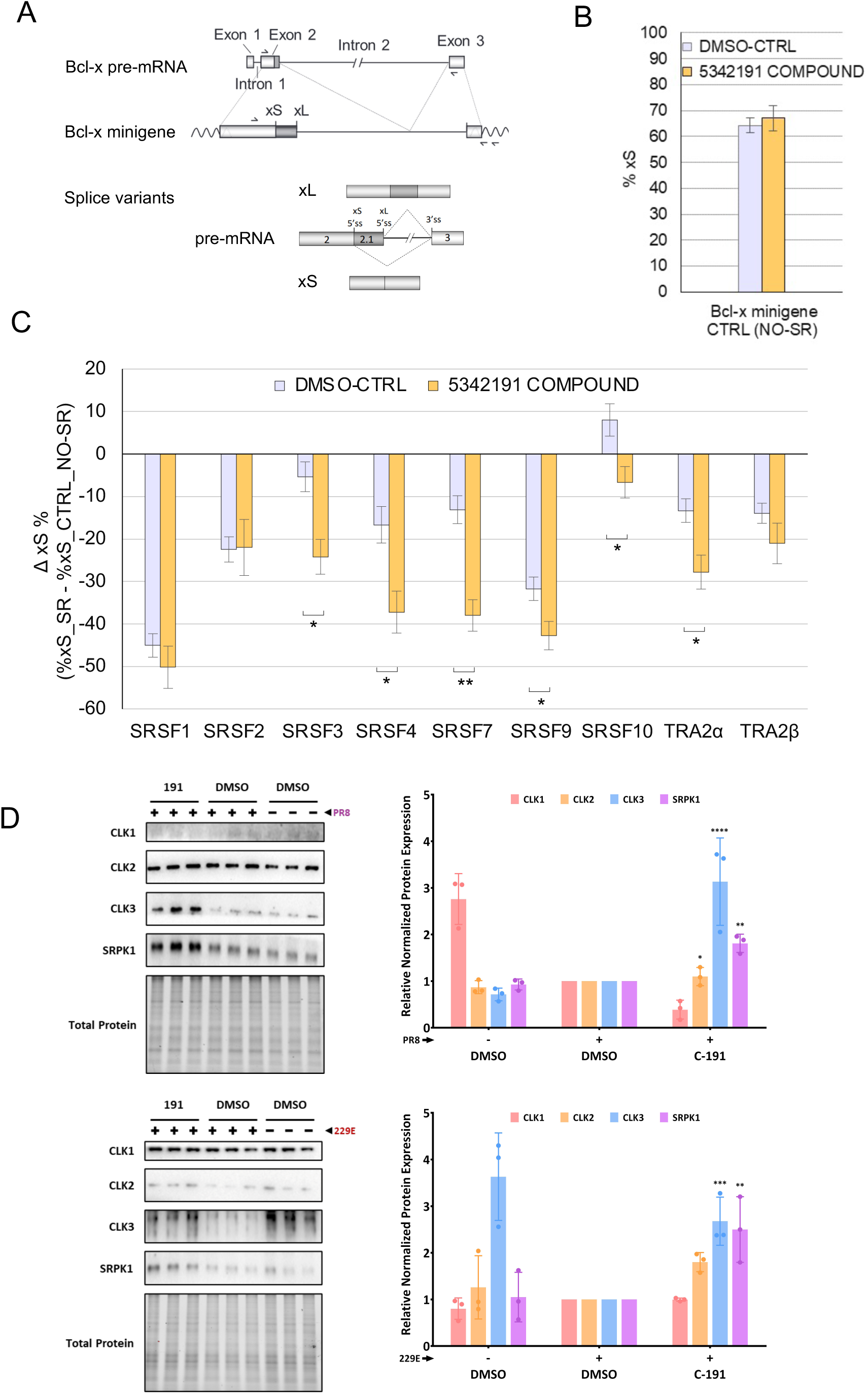
5342191 alters function of a subset of SR proteins/SR kinase expression. To measure the impact of 5342191 compound on the function of individual SR proteins, EcR293 cells were co-transfected with a *Bcl-x* minigene reporter plasmid and a plasmid expressing the indicated SR protein or an empty (No-SR) plasmid control. Six hours after transfection, EcR293 cells were treated with either 10 µM of 5342191 compound or only DMSO as control. Cells were harvested after 48 hours and total RNA was extracted. *Bcl-x* minigene-derived splicing products were amplified by RT-PCR using a minigene-specific pair of primers. [α-^32^P] dCTP was added to PCR mixture. Amplified products were fractionated onto a 4% native polyacrylamide gels that were scanned on a STORM PhosphorImager 860 to quantify the intensity of the bands using the ImageQuant software. The relative abundance of minigene derived Bcl-xS splice variant was expressed as percentage relative to the total abundance of both xS and xL splice variants. (A) Schematic representation of the human *Bcl-x* gene with relevant portions included in the minigene and derived splicing products. The positions of the primers used for RT-PCR assay is shown. (B) EcR293 cells co-transfected with the *Bcl-x* minigene and the empty No-SR Ctrl plasmid were treated for 48 hours with 10 µM of 5342191 compound or DMSO alone as control. The percentage of the minigene-derived xS splice variant is shown. (C) Changes in the relative abundance of the minigene-derived xS splice variant caused by expressing each SR protein in conditions of CTRL-DMSO or 10 µM 5342191 treated cells are plotted from the results of triplicate independent assays. Multiple unpaired t-tests were performed using GraphPad Prism version 10.2.3 and p values are represented by asterisks (*, p < 0.05, ** p < 0.01). (D) To assess the impact of 5342191 on SR kinase expression, Huh7 (-/+ HCoV-229E) or A549 (-/+ PR8) were treated with an EC_90_ dose of 5342191. 24 hpi, cells were harvested and proteins extracts analyzed by western blot to assess changes in expression of CLK1-3 and SRPK1. At the left, representative western blots and, right, a summary of changes in SR kinase expression over n>3 independent assays.

To understand how 5342191 affects SR protein function, we also examined its impact on the levels of various SR kinases known to regulate SR protein function through phosphorylation of their RS domains^26–28^. Western blot analysis of cell extracts from both A549 and Huh7 cells treated with DMSO or 5342191 (Fig. 13D) showed that the addition of 5342191 increased the levels of CLK3 and SRPK1 in both cell lines. Notably, viral infection also led to marked changes in the expression of select SR kinases, though the specific kinase affected varied between the viruses and cell lines: influenza PR8 infection of A549 cells resulted in reduced CLK1 levels while HCoV-229E infection of Huh7 cells led to a decrease in CLK3 abundance.

## Discussion

The development of broad-spectrum antibiotics has proven invaluable in controlling bacterial infections. However, the marked differences in sequence and replication strategies among viruses have complicated the development of similar broad-spectrum antivirals that target viral-encoded functions (i.e., viral protease, polymerase). Since viruses are obligate parasites, they rely heavily on host cellular functions for replication. Consequently, altering a cellular function necessary for the replication of multiple viruses offers the potential for developing a broad-spectrum antiviral. One such process is RNA processing, in particular splicing, which allows viruses to significantly expand the coding capacity of their genomes despite the size constraints imposed by the virus particle that mediates genomic material transfer between cells and hosts.

Recent studies have identified several small molecule inhibitors of HIV-1 replication (cardiotonic steroids, ABX-464, 1C8, GPS491, filgotinib, 791, harmine) that induce significant perturbations in viral RNA accumulation at doses that have limited effect on host RNA accumulation or processing ^9,11,12,29–34^. This observation raises the possibility that viruses are inherently more sensitive to small perturbations in cellular RNA processing due to their reliance on a specifc balance of the different spliced viral RNAs encoding factors essential for virus assembly. However, it remains unclear whether different viruses are sensitive to perturbations induced by the same compound; the demonstration that cardiotonic steroids, harmine, 1C8, and GPS491 inhibit the replication of several different viruses suggests that this is indeed possible ^11–13,30,35,36^.

Building on these findings, in this report we demonstrate that the benzoxadiazol-4-amine, compound 5342191, not only inhibits the replication of HIV-1 but also affects three unrelated viruses: adenovirus, coronaviruses, and influenza. In all instances, the compound acts post-entry as indicated by its ability to act after viral gene expression has initiated at concentrations similar or below those required to suppress HIV-1 replication (see Table 1)^14^. In the case of adenovirus, 5342191 addition at the end of the adsorption period had no effect on the transcript level of the first viral gene expressed, E1A, indicating that viral DNA delivery to the nucleus and initiation of transcription from the viral genome is unaffected. In contrast, expression of E1A-induced viral genes (E1B, E2A, E2B) is significantly reduced, indicating that E1A function is blocked. Although 5342191 has limited effect on E1A protein expression, it alters the processing of its transcript by preventing the accumulation of E1A 9S RNA (which encodes E1A 55R, not detected by the E1A antibody used). Since the time of of E1A 9S mRNA production correlates with the onset of E1B, E2A, and E2B RNA accumulation (16 hpi) and its product (E1A 55R) can induce expression of adenovirus early and late genes^37^, it is possible that preventing the shift in E1A RNA processing and expression of the 9S RNA encoded factor underlies the response observed. Adenovirus infection induces alterations in RNA splicing factors necessary for appropriate processing of viral RNAs, particularly through the dephosphorylation of SR proteins by a complex formed by adenovirus E4-orf4 and cellular PP2A phosphatase^38,39^. By preventing the expression of other viral genes^38–41^, 5342191 likely inhibits the reprogramming of the host splicing apparatus required for late gene expression.

**Table 1:** Effect of 5342191 on HIV-1/Coronavirus/influenza replication.

| **Influenza A Virus (PR8 )** |  |  |  |
| --- | --- | --- | --- |
| *TCID<sub>50</sub> Assay (Infectivity) – MOI of 2* |  |  |  |
| **Compound** | **EC<sub>50</sub>(μM)** | **EC<sub>90</sub>(μM)** | **CC<sub>50</sub>(μM)** |
| 5342191 | 0.075 ± 0.017 | 0.099 ± 0.008 | > 10 |
| Favipiravir | 1.2 ± 0.135 | 6.2 ± 0.332 | > 100 |
| Oseltamivir | 0.257 ± 0.040 | 1.5 ± 0.209 | > 100 |
| *qPCR Assay (Viral RNA Release) – MOI of 2* |  |  |  |
| **Compound** | **EC<sub>50</sub>(μM)** | **EC<sub>90</sub>(μM)** | **CC<sub>50</sub>(μM)** |
| 5342191 | 0.051 ± 0.042 | 0.104 ± 0.023 | > 10 |
| *qPCR Assay (Viral RNA Release) – MOI of 0.01* |  |  |  |
| **Compound** | **EC<sub>50</sub>(μM)** | **EC<sub>90</sub>(μM)** | **CC<sub>50</sub>(μM)** |
| 5342191 | 0.047 ± 0.013 | 0.149 ± 0.055 | > 1 |
| **Coronavirus (HCoV-229E )** |  |  |  |
| *qPCR Assay (Viral RNA Release) – MOI of 2* |  |  |  |
| **Compound** | **EC<sub>50</sub>(μM)** | **EC<sub>90</sub>(μM)** | **CC<sub>50</sub>(μM)** |
| 5342191 | 1.2 ± 0.196 | 2.5 ± 0.282 | 34 ± 0.755 |
| Boceprevir | 0.111 ± 0.024 | 1.2 ± 0.095 | > 100 |
| Molnupiravir | 0.025 ± 0.009 | 0.137 ± 0.023 | > 100 |
| *qPCR Assay (Viral RNA Release) – MOI of 0.01* |  |  |  |
| **Compound** | **EC<sub>50</sub>(μM)** | **EC<sub>90</sub>(μM)** | **CC<sub>50</sub>(μM)** |
| 5342191 | 0.073 ± 0.013 | 0.857 ± 0.056 | > 1 |
| **HIV-1** |  |  |  |
| **Compound** | **EC<sub>50</sub>(μM)** | **EC<sub>90</sub>(μM)** | **CC<sub>50</sub>(μM)** |
| 5342191 | 0.750 | 1.75 | > 10 |

Given the 5342191-induced shifts in HIV-1 and adenovirus RNA processing, it is not surprising that influenza replication was also impacted by 5342191 addition, as two of its segments (M and NS) undergo splicing to generate mRNAs encoding proteins (M2, NEP) critical for virion formation and release ^42,43^. siRNA screens have shown that influenza is sensitive to shifts in splice factor function, as depletion of CLK1, a kinase that phosphorylates SR proteins, or treatment with the CLK inhibitor TG003 results in loss of virus replication^44^. Consistent with 5342191 acting through changes in splice factor function, treatment of infected cells not only reduced viral RNA accumulation but also affected the accumulation of viral spliced RNA (M2 and NS2). However, addition of 5342191 impacts the production of all viral RNAs as indicated by the loss of EU-labelled viral RNAs within 4 h of compound addition to the culture (Fig. 10), indicating that the major effect of this compound is on viral polymerase function.

In contrast to HIV-1, adenovirus, and influenza, whose RNAs undergo splicing, the entire replication cycle of coronaviruses occurs in the cytoplasm, and there is no evidence that any of their RNAs are spliced^19,22^. However, studies have identified several SR kinase inhibitors (SRPIN340, harmine) that can block coronavirus replication^12,45–48^. Such effects are explained in part by the presence of arginine-serine (RS) repeats within the viral nucleocapsid (N), with phosphorylation by SR kinases impacting N function within the viral RNA replicase complex^45–48^. As detailed in this study, 5342191 inhibited a range of coronaviruses including members of the alpha (HCoV-229E) and beta (HCoV-OC43, SARS-CoV2) coronavirus families,suggesting that it affects a process essential to all of them. Inhibition of both HCoV-OC43 and SARS-CoV-2 in human lung organoids indicates that 5342191’s antiviral properties are not limited to transformed cell lines. Analyses indicate that 5342191 does not affect the stability of the viral replicase complexes once formed (as indicated by the persistence of dsRNA and viral genomic RNA staining in perinuclear structures, Fig. 7), but rather alters viral polymerase function. Coronavirus polymerase generates multiple viral RNAs using a nested RNA strategy involving discontinuous copying of the genomic RNA^19,21^. The preferential reduction in nested versus genomic RNA accumulation after delayed addition of 5342191 (Fig. 6) suggests that the compound induces changes favoring continuous rather than discontinuous transcription by the viral polymerase complex, similar to that observed upon harmine addition^12^.

The ability of 5342191 to inhibit a range of viruses raised the question whether its use would confer additional advantages over existing direct-acting antivirals (DAAs). One potential advantage could be a greater barrier to the development of drug-resistant viruses. Consistent with this hypothesis, we observed (Fig. 11) that only 4 passages of either HCoV-229E or influenza PR8 in the presence of EC_90_ concentration of DAAs was sufficient to select for variants able to replicate with similar efficiency in the presence or absence of the drugs. In the case of HCoV-229E, the virus selected in the presence of 5342191 replicates to lower levels compared to the initial strain, and viral RNA accumulation in the media was further diminished upon addition of the compound (after 4 or 10 serial passages, Fig. 11B). In the case of influenza PR8, all compounds selected viruses with enhanced release of viral RNA into the media. However, unlike favipiravir or oseltamivir, the virus selected with 5342191 remained sensitive to compound addition after 4 rounds of selection. Together, these observations suggest a higher barrier to the development of resistance. Notably, the EC_90_ doses of 5342191 required to suppress virus replication are well below the CC_50_ and doses previously shown to have minimal effect on host gene and protein expression^14^. Further studies in mice (Table S2-4) determined that 5342191 is orally bioavailable and well tolerated at a daily oral dose of 40 mg/kg over 7 days. Using an oral dose of 30 mg/kg, a blood concentration of 2 µg/ml (5.5 µM) was maintained up to 8 h after administration. The absence of significant changes in body weight or abnormal findings upon gross necroscopy of 5342191-treated animals indicate that the compound is well tolerated.

Investigation of how 5342191 achieves its antiviral effect on coronaviruses and influenza revealed a response similar to that observed previously in the context of HIV-1 ^14^. The ability of inhibitors of G protein coupled receptors (GPCRs) and MAPK signaling pathways to restore coronavirus and influenza virus replication in the presence of 5342191 suggests that the compound is an activator of a GPCR that signals through MAPK activation to alter splice factor activity. The alterations in adenovirus/influenza RNA splicing, coupled with the similarity in the response of harmine^12^, an SR kinase inhibitor, to coronaviruses suggested that 5342191 may work through the effects on SR protein/SR kinase function. Previously, we reported that 5342191 treatment altered accumulation of select SR proteins, with a decrease in levels of SRSF1 and SRSF3 and an increase in SRSF4^14^. However, RNA-seq data analysis of treated cells did not reveal a change in abundance of the mRNAs for these factors, indicating that regulation must be occurring at the post-transcriptional level^14^. In this report, we determined that 5342191-treatment also alters the activity of select SR proteins in the context of *Bcl-x* RNA splicing, significantly enhancing the activity of SRSF3, SRSF4, SRSF7, SRSF9, and Tra2a, while reversing the activity of SRSF10. In the absence of altered abundance of mRNAs, changes in SR protein function may be the results of the observed changes in expression of select SR kinases when cells are treated with 5342191. Previous studies have highlighted the important role of SR kinases in the replication of multiple viruses^49,50^ and how modulation in their relative expression can impact virus gene expression ^51^. Furthermore, cell signaling is known to impact SR protein/SR kinase function ^52–54^. Our observation that 5342191 treatment of Huh7 or A549 cells selectively increased levels of both CLK3 and SRPK1-kinases known to phosphorylate host SR proteins^55,56^ and coronavirus N proteins^46–48^-is consistent with the hypothesis that 5342191 suppresses virus replication through induction of signaling events to affect SR kinase expression.

Taken together, our observations demonstrate the potential of targeting cellular processes essential for the replication of multiple viruses, including several of current concern (HIV-1, coronaviruses, influenza), and that virus suppression can be achieved at doses with minimal effect on cell/mouse viability. The broad-spectrum anti-viral activity of 5342191 coupled with the higher barrier to the emergence of resistant virus strains underline the strengths of such an approach over DAAs. Moving forward, we plan to pursue medicinal chemistry studies focused on identifying 5342191 derivatives with improved antiviral activity and therapeutic index (CC_50_/EC_50_), which could be useful in virus challenge studies in animals, a critical step toward unlocking the therapeutic potential of this strategy.

---

**Table S1:**
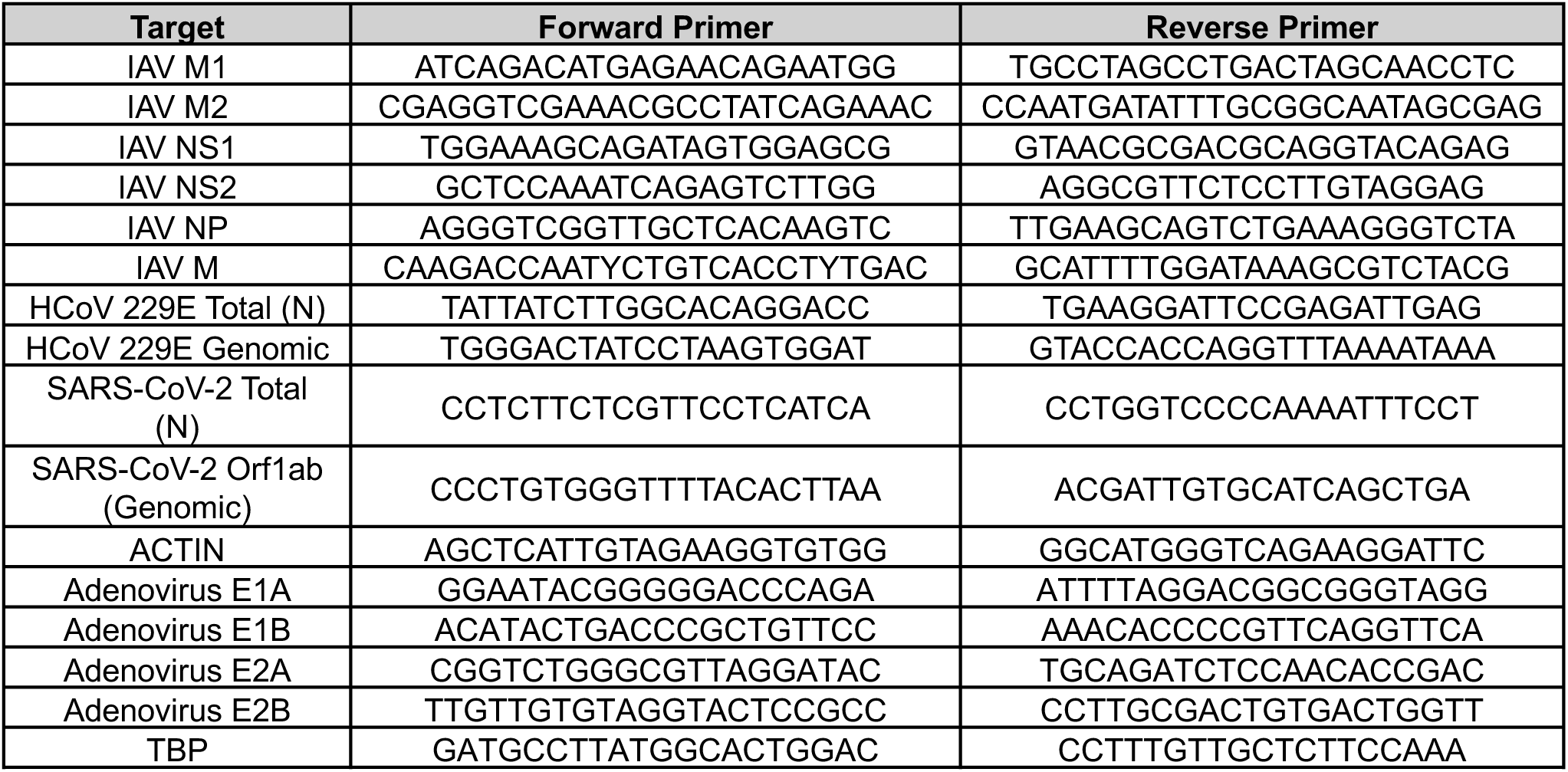
Sequences of RTqPCR primers.

**Table S2:** Summary of Body Weight Changes After Repeat Daily Oral 5342191 (C-191) Treatment.

| Treatment | Dose (mg/kg) | Mean Body Weight ± SD (g) | | Mean Body Weight Changes Day 1-8 ± SD (g) | Animal number |
| --- | --- | --- | --- | --- | --- |
|  |  | Day 1 | Day 8 |  |  |
| Vehicle Control | 0 | 24.7 ± 1.6 | 26.1 ± 2.0 | 1.4 ± 0.7 | 4 |
| C-191 | 50 | 26.6 ± 3.7 | 27.7 ± 5.8 | 1.1 ± 0.6 | 2 |
| | 40 | 28.0 ± 1.7 | 27.6 ± 1.7 | -0.4 ± 0.7 | 4 |
| | 30 | 26.1 ± 1.3 | 26.1 ± 0.9 | 0.1 ± 0.5 | 4 |
| | 10 | 27.8 ± 2.0 | 28.1 ± 1.8 | 0.4 ± 0.2 | 4 |

**Table S3:**
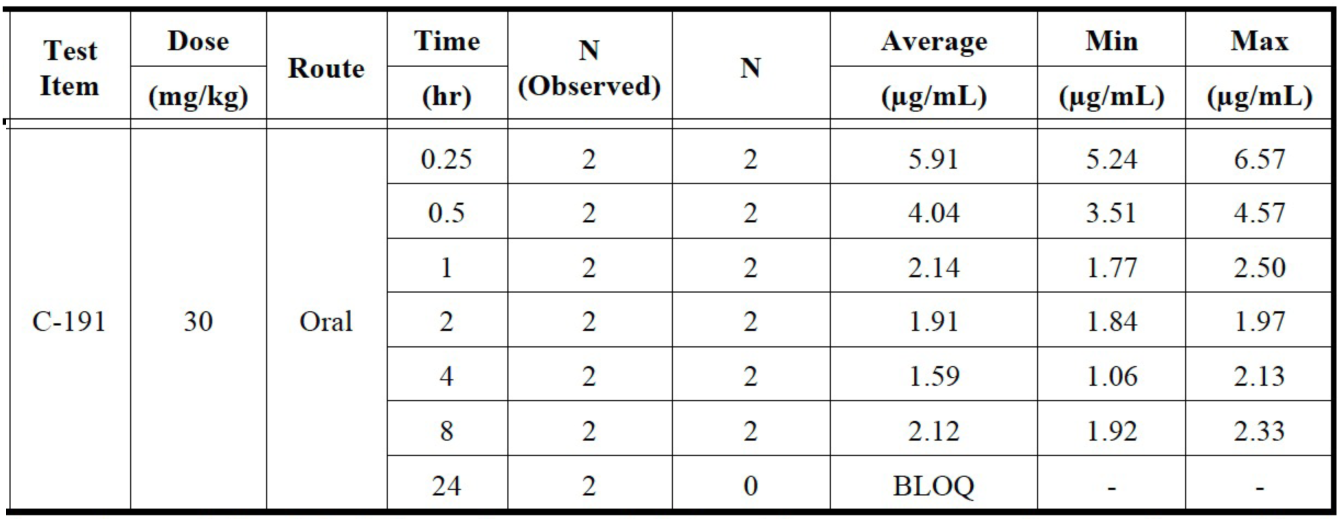
Mean Plasma Concentrations of Oral Dose of 5342191 (C-191) in Female Mice.

**Table S4:** Mean Group PK Parameters of Oral Dose 5342191 (C-191) in Female Mice.

| Test Item | Dose (mg/kg) | Route | C<sub>max</sub> (μg/mL) | T<sub>max</sub> (hr) | AUC<sub>last</sub> (hr*μg/mL) | AUC<sub>INF</sub> (hr*μg/mL) | AUC<sub>Extrap</sub> (%) | V<sub>z</sub>/F (mg/(μg/mL)/kg) | Cl/F (mg/(hr*μg/mL)/kg) | MRT<sub>last</sub> (hr) | K<sub>el</sub> (1/hr) | T<sub>1/2</sub> (hr) | R<sup>2</sup> |
| --- | --- | --- | --- | --- | --- | --- | --- | --- | --- | --- | --- | --- | --- |
| C-191 | 30 | Oral | 5.91 | 0.25 | 16.48 | 59.70 | 72 | 10.23 | 0.50 | 3.73 | 0.050* | 14.11* | 0.18* |

## Notes

### Competing Interest Statement

The authors have declared no competing interest.

